# Rational reduction of a sorghum SynCom that preserves growth promotion reveals flavonoid-mediated plant microbe interactions

**DOI:** 10.64898/2026.03.20.709941

**Authors:** Dean Pettinga, Citlali Fonseca-García, Genevieve Krause, Hannah Ploemacher, Travis Wheeler, Chaevien S. Clendinen, Pubudu Handakumbura, Robert Egbert, Devin Coleman-Derr

## Abstract

- Plant growth is influenced by the composition of its associated microbiome. The inherent complexity and functional redundancy of natural plant microbiomes presents a formidable barrier to understanding the myriad biological interactions therein. Efforts have been made to develop synthetic microbial communities (SynComs) that can provide a rigorous and generalizable framework for the rational design of next-generation microbial products for sustainable agriculture. We test multiple strategies for stable, plant growth promoting SynCom design and evaluate the phenotypic and molecular impacts of a successful plant-SynCom interaction.
- We designed 4 distinct, reduced-complexity variants of SynCom SRC1 and assessed their capacities for colonization, stability, and plant growth promotion. To understand the impact on plant performance of our highest performing SynCom variant, we characterized the host’s longitudinal transcriptional response to SynCom inoculation and corroborated the results with metabolomics analysis.
- The top performing SynCom stably colonized sorghum roots and rhizospheres, elicited plant growth promotion, and induced dynamic spatiotemporal gene transcription in sorghum roots and shoots defined by modulation of growth-defense tradeoff machinery and enhanced flavonoid production.
- The resultant reduced-complexity SynCom is a highly stable, soil-independent, plant growth promoting, and demonstrates the utility of colonization-based selection criteria, integrated with longitudinal transcriptomic and metabolomic characterization.

## Introduction

The microbial communities associated with roots – the rhizosphere and endosphere – are highly diverse and functionally complex, representing a ’black box’ of interactions that govern host health and nutrient access (Philippot *et al*., 2013; Trivedi *et al*., 2020). While large-scale metagenomic studies have established strong correlations between microbial diversity and plant fitness, the sheer complexity of natural microbiomes makes identifying causal relationships a significant challenge (Mueller & Sachs, 2015; Levy *et al*., 2017). Synthetic Microbial Communities (SynComs), small subsets of isolates representing natural communities that retain the full set of desired functional outputs are emerging as essential tools in microbiome science because they enable researchers to reduce this complexity to a defined, tractable system (Xu *et al*., 2025). By selecting and combining specific, culturable isolates, SynComs allow for controlled ecological experiments to test hypotheses about inter-species interactions, competition, resource sharing, and the molecular mechanisms that facilitate beneficial host-microbe relationships (Martins *et al*., 2023). Thus, SynComs shift the field from broad ecological observation toward precise, modular, and permutable experimental systems, enabling systematic dissection of the fundamental rules governing microbiome assembly and function.

Beyond basic research, one major translational goal of SynCom design is to enable the development of cost-effective, robust, and scalable microbial bio-inoculants for sustainable agriculture (Díaz-Rodríguez *et al*., 2025). Translating beneficial SynComs from the lab to the field presents unique engineering challenges. Complex communities often prove unstable under fluctuating environmental conditions (Finkel *et al*., 2017; de Souza *et al*., 2019, 2020) and the synthesis effort scales with the number of members. Success in agricultural application therefore hinges on identifying a minimal, stable SynCom. This pursuit requires a systematic approach to community reduction, stability testing across host compartments, and, critically, understanding the molecular dialogue that ensures SynCom persistence and efficacy within the host system.

Our foundation for this work stems from a previously reported beneficial synthetic community named SRC1 (Fonseca-García *et al*., 2024). This SynCom, comprising 57 bacterial isolates originally derived from the rhizosphere of field-grown *Sorghum bicolor*, demonstrated robust host growth. *Sorghum bicolor* is a globally important crop known for its abiotic resilience and ability to thrive in marginal lands (Calviño & Messing, 2012; Brenton *et al*., 2016; Appiah-Nkansah *et al*., 2019), making it an ideal target for microbiome engineering efforts. While the SRC1 community proved highly effective at colonization and growth promotion, its relatively large size proved challenging for regular deployment in the context of laboratory research, and would additionally present a logistical and economic bottleneck for commercial application. This established the primary goal for the current study: to systematically leverage the functional data gathered for the SRC1 members to design a community with a greatly reduced number of isolates, and validate that it is both stable and functionally equivalent to the parental SRC1 community.

In the present study, we tackled this challenge by formulating four distinct, second-generation communities – the SRC2 series – using rational selection principles aimed at maximizing either: 1) phylogenetic diversity, 2) predicted genomic functions, 3) positive host benefit associated interactions, or 4) robust root colonization. We assessed these SRC2 SynComs for stability (alpha- and beta-diversity) and their capacity to increase sorghum shoot biomass. The community selected to maximize root colonization, referred to as SRC2v4, was the top-performing, reduced complexity community that successfully retained many of the host benefits of SRC1. To uncover the mechanisms underpinning this successful stability and functional retention, we conducted an integrated analysis of the host’s transcriptional and metabolic response to SRC2v4. We identified dynamic activation of the host’s flavonoid biosynthetic pathway in response to SRC2v4, which led to the robust accumulation of flavonoid metabolites in the root tissue. This suggests that the sorghum host actively accommodates the SRC2v4 community by modulating secondary metabolism. Together this work explores rational redesign and evaluation of a SynCom and demonstrates complex interactions underpinning the beneficial interactions leading to strong colonization and increased plant performance.

## Materials and Methods

### SynCom Designs

Four distinct strategies were employed to reduce the complexity of SRC1 (Fonseca-García *et al*., 2024), a synthetic bacterial community of 57 members, down to 15 members (**Fig. 1, S1, Supplement SD1**). For all SRC2 variants (v1-v4) formulations, we ensured that the 15 selected isolates did not share identical 16S rRNA gene sequences in the V3-V4 region in order to maximize our ability to resolve abundance patterns for each member of the community.

**Figure 1.**
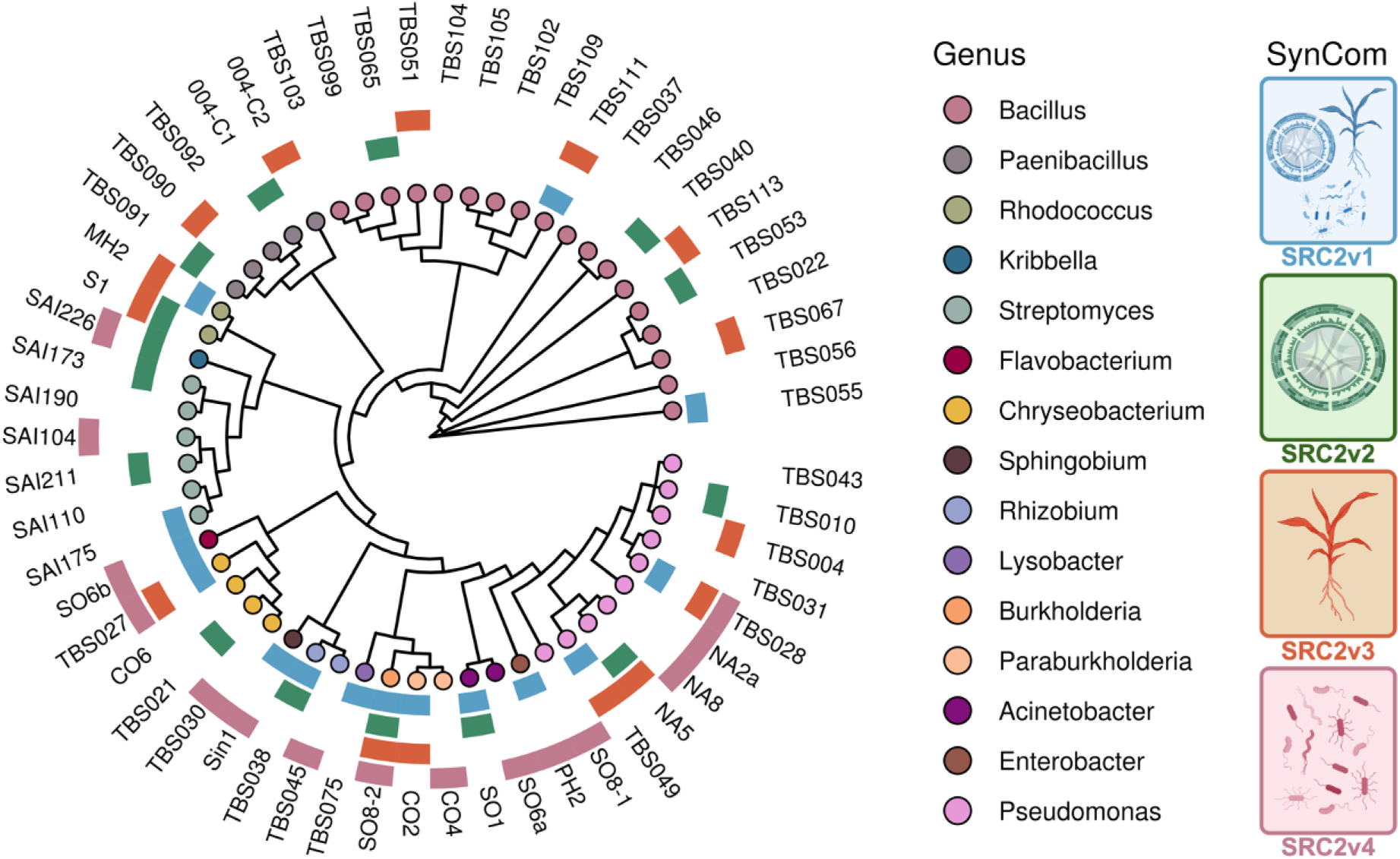
Synthetic community complexity reduction designs. Phylogenomic cladogram of SRC1 members built using multiple alignment of 74 single copy bacterial genes using *GToTree* (Lee, 2019). Tree tips are colored by Genus. Outer rings identify membership of each isolate in different reduced-complexity SynCom variants, colored in accordance with the pictogram at right.

The first variant, named SRC2v1, was designed to closely reflect the taxonomic composition of SRC1, maximizing phylogenetic diversity. Unique isolates from 11 of the 17 genera in SRC1 were selected for SRC2v1. Two isolates each were selected for the most abundant genera in SRC1: *Bacillus* (18/57) and *Pseudomonas* (9/57). In total, 15 isolates from 13 genera were selected for SRC2v1.

Second, SRC2v2 was designed to maximize predicted functional capacity. We used *OrthoFinder* (Emms & Kelly, 2015, 2019) to identify protein orthogroups across all SRC1 genomes, where an orthogroup is defined as a set of genes descended from a single gene in the last common ancestor of the considered genomes. Because many orthogroups lack functional annotation and interactions between orthogroups are not generally understood, we adopted the simplistic goal of maximizing the total collection of orthogroups retained across the entire community. Specifically, we sought to reduce the number of genomes while maximizing the number of orthogroups that are found in at least one of the remaining genomes. This is a variant of the NP-hard Set Cover problem (Karp, 1972), so we implemented a simple greedy strategy of genome removal: (phase 1) letting C_i_ be the number of remaining genomes that include orthogroup i, repeatedly remove a genome that maximizes the smallest value of C_i_ among remaining genomes, repeating until enough genomes have been removed to force C_i_=0 for some orthogroup i. (phase 2) Repeatedly remove a genome that minimizes the number of orthogroups i that are set to C_i_=0 as a result of the removal. This removal process continued until 15 genomes remained; heuristically selecting a set of 15 isolates that collectively contain as many orthogroups (and therefore as much potential function) as possible.

Third, SRC2v3 was designed using data from a permutational membership experiment based on the original SRC1 design (data unpublished). In this experiment, we systematically and separately depleted each of the three most abundant genera in SRC1 (*Bacillus*, *Pseudomonas*, or *Streptomyces)* to assess their individual contribution to plant phenotypes. Using 16S rRNA amplicon sequencing data and plant phenotypic measurements from this experiment, we evaluated the relationship between shoot growth and bacterial abundance for each member. Based on these analyses, strains exhibiting the strongest positive associations with shoot biomass were selected for inclusion in SRC2v3 (**Supplement SD2**). The permutational analysis further highlighted an important generalizable role for *Bacillus* and *Pseudomonas* members in promoting positive plant phenotypes; therefore, strains belonging to these genera were prioritized during formulation.

Lastly, SRC2v4 was formulated to maximize colonization strength. Using relative abundance data derived from the same permutational membership experiment, we ranked strains by their cumulative relative abundance (as measured by 16S rRNA amplicon data) across all root samples and selected 15 of the most abundant members (**Supplement SD3**). Among strains with highly similar relative abundance, priority was given to strains previously identified as being capable of utilizing sorghum root exudates as a carbon source (Fonseca-García *et al*., 2024).

### Microbox Experimental Setup

To investigate *in planta* colonization and the effects of sorghum root consortium (SRC1) strains on plant phenotypes, a series of controlled experiments were conducted using *Sorghum bicolor* cultivar RTx430. Seeds were surface-sterilized in 10% (v/v) sodium hypochlorite for 15 minutes, rinsed five times with sterile distilled water, and germinated on sterile, moistened filter paper at 30°C in the dark.

To create the SRC2 SynCom variants, bacterial strains were grown on tryptic soy agar (TSA; 15 g pancreatic digest of casein, 5 g peptic digest of soybean meal, 5 g NaCl, and 15 g agar per liter) at 30°C according to their growth rate (Fonseca-García *et al*., 2024). For each strain, six inoculating loopfuls containing a mixture of cells, exopolysaccharides, and other bacteria-derived biomass were resuspended in 1.2 mL sterile 1× PBS (1 loop ∼12 mg biomass per 200 uL of 1X PBS). Cell suspensions were evenly pooled according to the desired consortium composition and aliquotted into 50 mL sterile beakers containing 6 mL per inoculum. For heat-killed (HK) treatments, pooled suspensions were autoclaved once at 120°C for 25 minutes prior to use.

Seedling assays were conducted in 5 L sterile microboxes (Combiness, Belgium) containing 1 kg of autoclaved calcined clay (Sierra Pacific Supply, USA) mixed with 400 mL of sterile Hoagland’s solution (1.6 g L⁻¹; Sigma). Pre-germinated seeds were incubated for 10 min in the respective inoculum (SRC2v4, HK, or Mock [1× PBS]) before planting 5 cm below the surface, followed by an additional 1 mL inoculum applied to each seed (four germinated seeds per box). Growth conditions were maintained at 26–28°C, 16 h:8 h light:dark, and 50% relative humidity. Plants were watered with 6 mL of sterile distilled water per plant once a week. After one week, the lids were removed in a laminar flow hood and a second inverted box was sealed to the container as an alternative lid, providing additional room for foliar growth. At harvest (1 month post-inoculation), shoots were collected for phenotyping and biomass measurements, while roots were used for DNA and RNA extraction and/or metabolite profiling, depending on the experiment. Throughout all experiments, individual plants within each microbox were treated as independent biological replicates, as root systems were physically separated within the growth substrate and sampled individually. The above growth conditions and preparations were applied to each the following experimental designs, with the following modifications:

*1) SRC2v1–v4* (**Fig. 2-3, S1-S5**): Each version of the SRC2 corresponded to iterative reduced variants of the SRC1 root bacterial community, differing in strain composition (**Supplement SD1**). These experiments followed the general setup described above; each treatment included a minimum of four replicate microboxes, each containing four plants which were measured for shoot fresh and dry weight, shoot length, and water content. Metagenomic DNA was collected from inocula, roots, rhizospheres, and soils 16S rRNA analysis.
*2) SRC2v4 vs. HK (heat-killed)* (**Fig. 4a**): To assess the contribution of microbial activity, plants were inoculated with either live SRC2v4 or HK inocula (autoclaved at 120℃ for 25 min). Growth and harvest procedures were as described above and shoot fresh and dry weight, shoot length, and root fresh weight were measured.
*3) SRC2v4 vs. SRC1* (**Fig. 4b**): Comparative inoculations were performed using live SRC2v4 and the previously established SRC1 consortium under identical growth conditions (Fonseca-García *et al*., 2024). A similar amount of cell mass was added from both formulations for treatment. Host phenotypes were measured including: shoot fresh and dry weight, shoot length, and root fresh weight.
*4) Metatranscriptomics* (**Fig. S6**): To assess microbial transcriptional activity in the rhizosphere, plants were grown as described above and inoculated with live SRC2v4. Rhizosphere samples were collected 1 month post-inoculation and two replicate microboxes per treatment were sampled. Collected rhizosphere material was immediately flash-frozen in liquid nitrogen and stored at -80℃ prior to RNA extraction and sequencing.
*5) Time-course RNAseq* (**Fig. 5-6, S7-S10**): For transcriptomic profiling, plants inoculated with live SRC2v4 or HK inocula, or with a Mock treatment, were harvested at multiple time points post-inoculation (1 h, 24 h, 72 h, 1 week, 2 weeks, and 1 month) under the same growth and harvest conditions described above. SRC2v4 and Mock were harvested at all time points, while HK was harvested at 24 h, 1 week, and 1 month post-inoculation. At each time point, shoot and root tissues were harvested at 12:30 PM, within a 30-minute harvest window, and immediately flash-frozen in liquid nitrogen and stored at -80℃ until RNA and metagenomic DNA extraction.
*6) Metabolomics* (**Fig. 7, S11**): To characterize metabolite profiles associated with SRC2v4, plants were grown as described above, and plants inoculated with live SRC2v4 or Mock treatment were harvested 1 month post-inoculation. Harvests were performed at 12:30 PM within a 30-minute collection window. Each treatment included four replicate boxes, each containing four plants. Roots and rhizosphere material were collected for downstream metagenomic DNA, metabolite extraction, and LC-MS analysis (described below).

**Figure 2.**
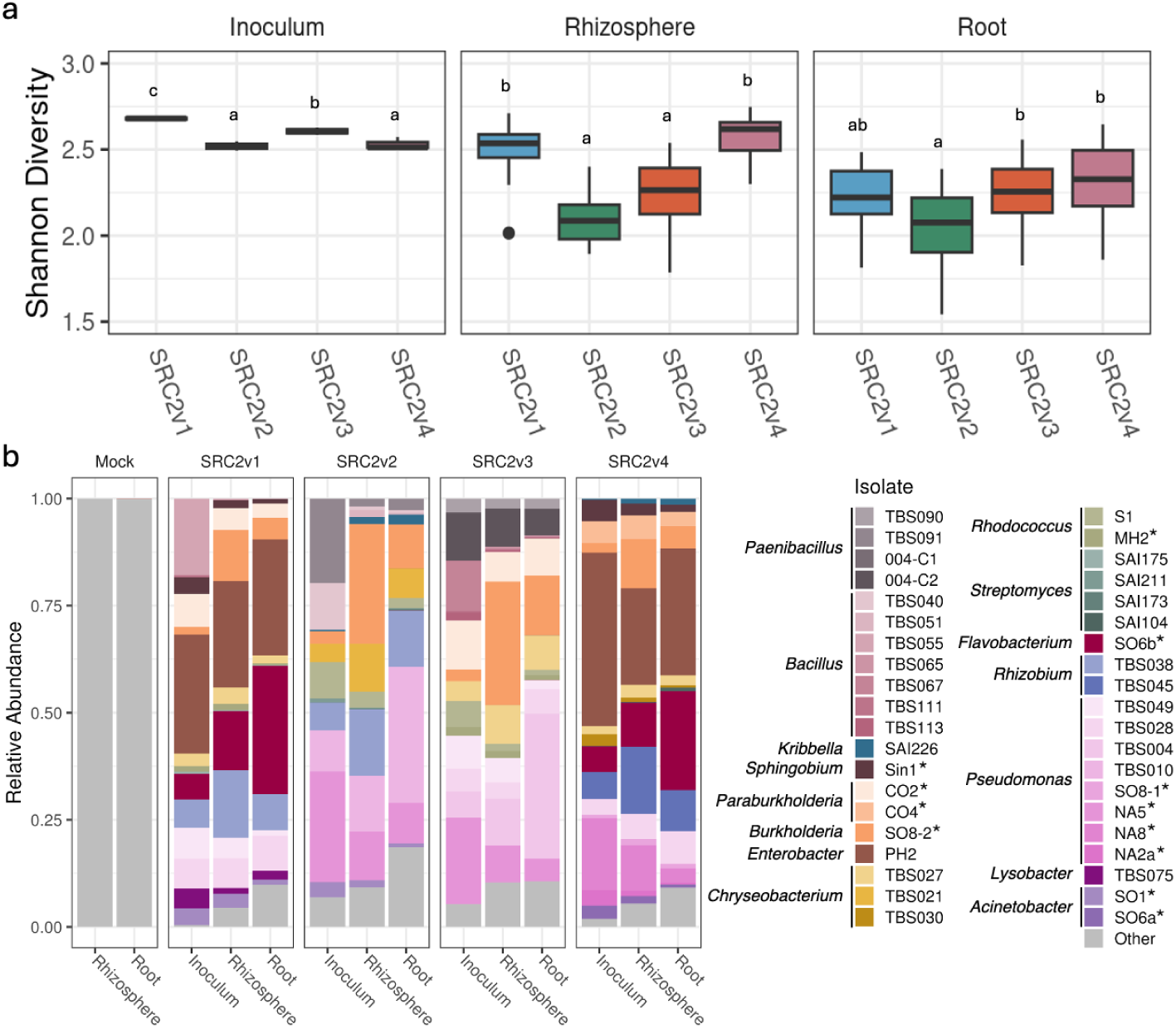
Reduced-complexity SynComs SRC2v1-v4 exhibit variable stabilities across plant compartments. **a)** Shannon Index across colonization compartments illustrates variable filtering effects of host plant on diversity between SynCom treatments. Diversity comparisons of SRC2v1-4 analyzed within compartment: inoculum, rhizosphere, and root. In all panels, letters above the boxplots represent statistical significance based on one-way ANOVA followed by post-hoc Tukey’s HSD (alpha = 0.05). **b)** Mean relative abundances of each SynCom illustrate community composition across compartments. (*sorghum exudate metabolizers, see **Supplement SD1**)

**Figure 3.**
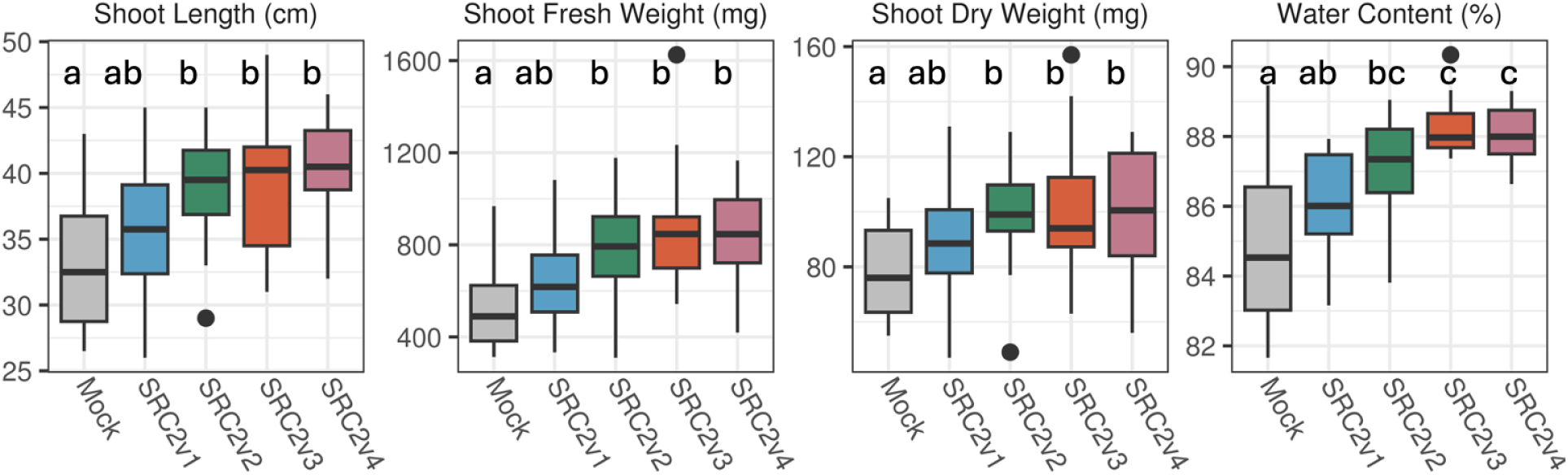
SynCom variants induce variable impacts on sorghum host phenotypes. SRC2v1 did not significantly differ from Mock treatment, but SRC2v2, SRC2v3, and SRC2v4 all increased shoot length, shoot fresh weight, shoot dry weight, and water content relative to Mock (ANOVA one-way, Tukey post hoc test adjusted p-value <0.05).

**Figure 4.**
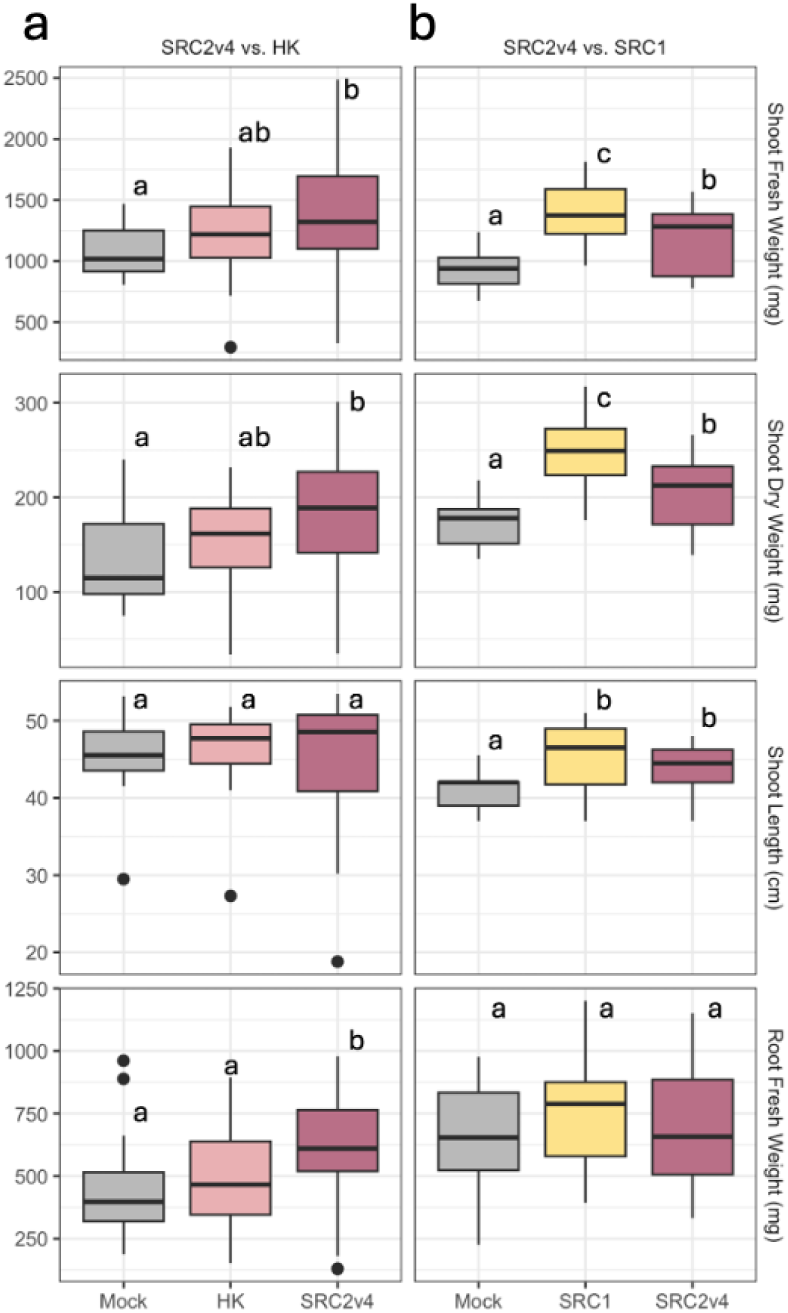
Host plant growth benefits upon treatment with live SRC2v4 treatment benchmarked against both HK **a)** and larger, 57-member SRC1 community **b)** in two separate experiments. Surface-sterilized, germinating seedlings were inoculated with each treatment, planted in sterilized 5L microboxes with 1kg of calcined clay. After one month of growth, samples from each treatment were harvested and plant host phenotypes were measured. Letters represent significant differences (p<0.05), ANOVA with post-hoc pairwise Tukey HSD.

**Figure 5.**
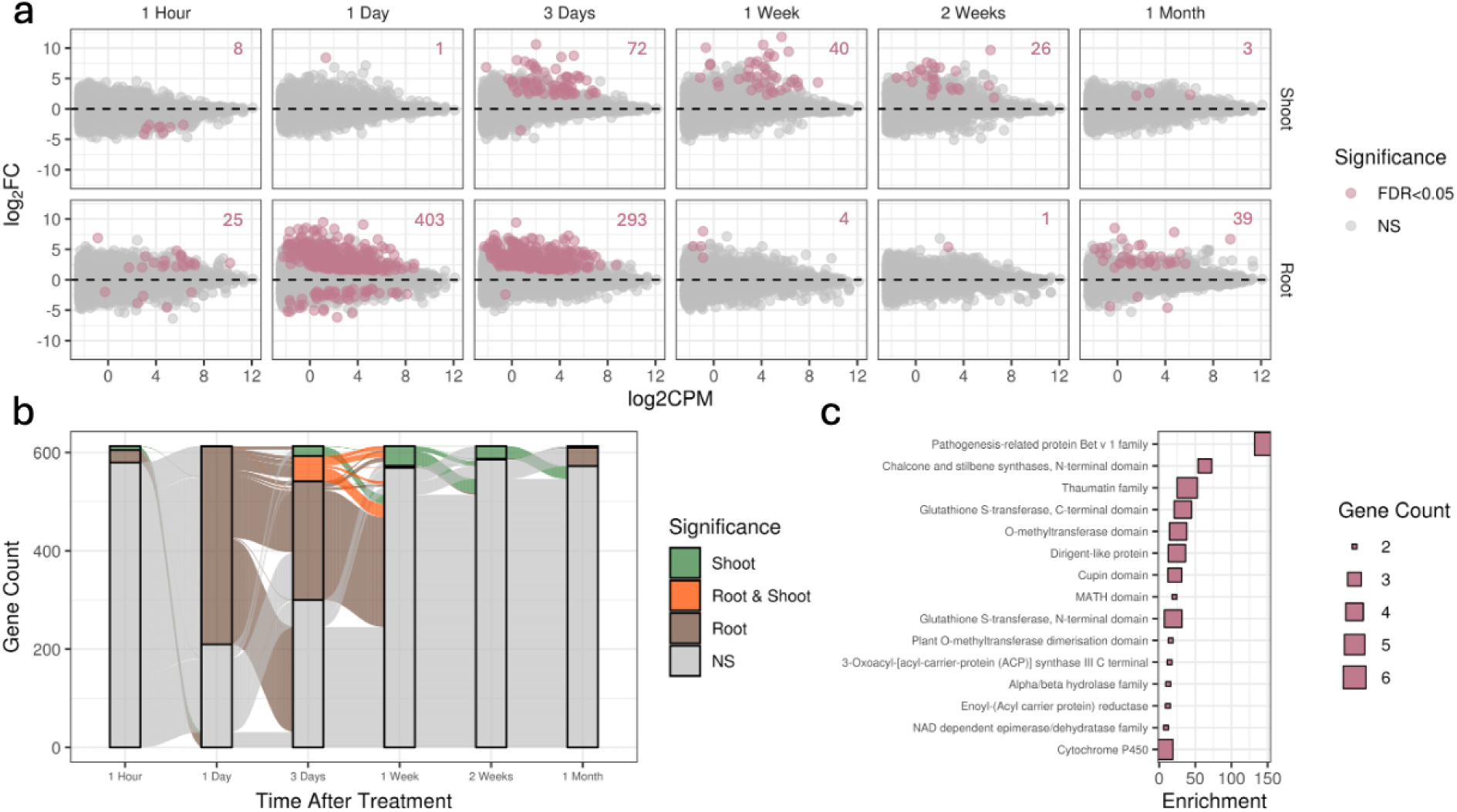
Sorghum host plants respond to SRC2v4 treatment with dynamic, spatiotemporal transcriptomic regulation. **a)** Differentially expressed (DE) genes responding to SRC2v4 treatment relative to Mock across six time points after treatment: 1 hour, 1 day, 3 days, 1 week, 2 weeks, and 1 month in both root (lower) and shoot (upper) tissue. Numbers in the upper right of each panel represent DE gene counts. Log_2_FC: log_2_-fold change (SRC2v4/Mock), logCPM: average log_2_counts per million across SRC2v4 & Mock treatments, FDR<0.05: significantly differentially expressed genes after multiple testing correction, NS: not significant. **b)** Significantly differentially expressed genes in response to SRC2v4 across the experiment reveals early regulation of genes in root or shoot that become differentially regulated in other tissues at later timepoints. Total gene count represented by y-axis and timepoints on x-axis. Genes are colored by tissue in which significant differential expression occurs. **c)** Enriched PFAMs among the genes initially regulated in root tissue, then in shoot tissue at a later timepoint during the experiment (n=73). Presented terms represent significant hypergeometric enrichment results in this subset relative to all genes in the genome after multiple testing correction. Enrichment=(PFAM gene count/selection gene count)/(PFAM genes in genome/genome gene count).

**Figure 6.**
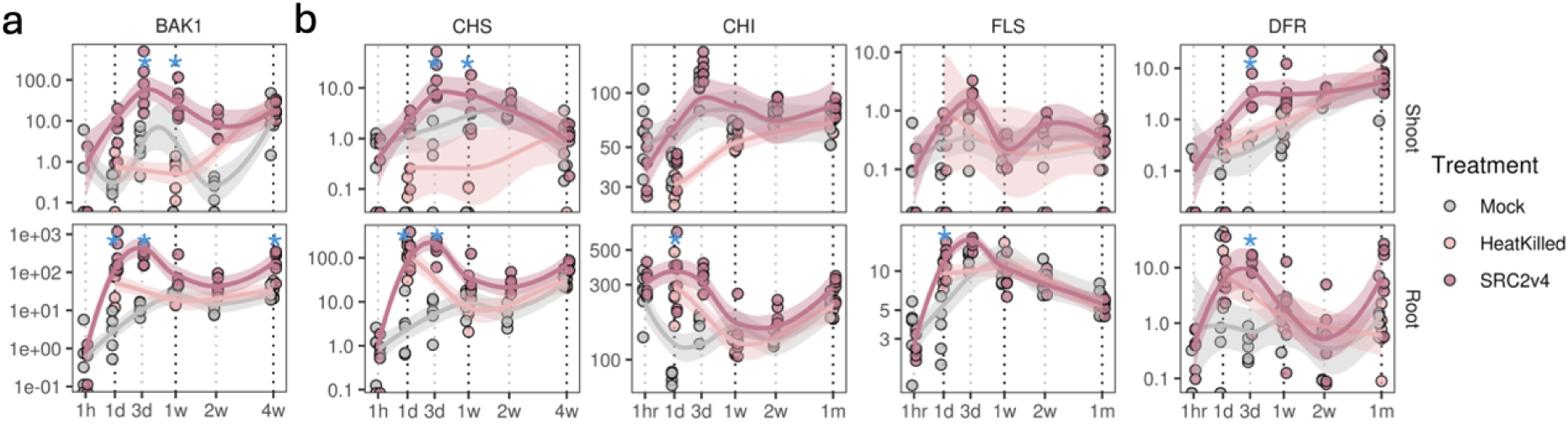
Spatiotemporal transcriptional regulation of key genes in SRC2v4 perception and plant response. Each gene is presented in two panels with treatment comparisons within each tissue type shoot (upper) and root (lower) across all time points with Loess fit trendlines by treatment. **a)** BAK1 microbial and brassinosteroid perception receptor like kinase exemplifies spatiotemporal DE patterns observed across the genome **b)** Flavonoid biosynthetic genes CHS, CHI, FLS, and DFR display significant upregulation upon SRC2v4 treatment. **c)** Hormone modulators include Nodulin, CK-N-Gluc, CK-O-Gluc, and MES. Blue stars indicate significant upregulation in SRC2v4 relative to Mock (Log_2_FoldChange > 1, FDR < 0.05), CPM = counts per million reads.

**Figure 7.**
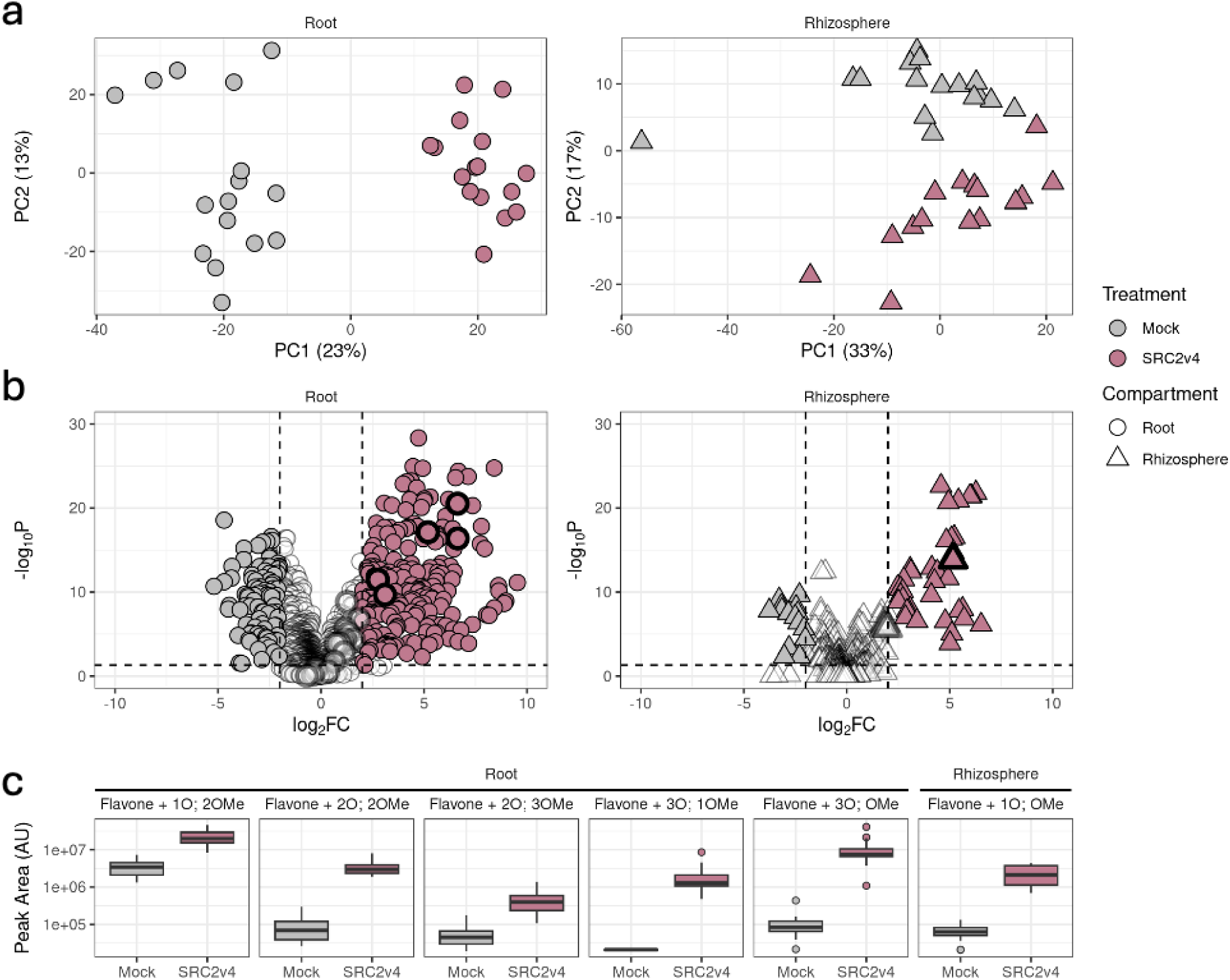
LC-MS of roots and rhizospheres inoculated with SRC2v4 reveals significant differentiation of metabolome. **a)** Principal Component Analysis of root (left) and rhizosphere (right) exhibits significant differentiation by treatment in root (PERMANOVA: Root R^2^ = 0.22, p = 0.001; Rhizosphere R^2^ = 0.16, p = 0.001). **b)** Volcano plots of differentially abundant metabolites colored by significance: white = not significant, pink = up in SRC2v4, gray = up in Mock; Log_2_FC = log_2_FoldChange(SRC2v4/Mock). Highlighted points with wide outlines are annotated flavonoid compounds. **c)** Significantly enriched flavonoid compounds grouped by compartment of enrichment, and colored by treatment.

### 16S rRNA Amplicon Metagenomics

Extraction of both rhizosphere and root compartment (including both rhizoplane and endosphere bacterial DNA) was performed using the Qiagen DNeasy PowerSoil DNA Isolation Kit (Cat. # 12888-100; QIAGEN, Venlo, Netherlands) with MoBio Power Soil DNA isolation kit (Catalog No. 12888-100; MoBio, Carlsbad, CA, USA) with 150 mg of flash-frozen root tissue or 200 mg of rhizosphere ground at 25 Hz for 5 min using Qiagen Tissue Lyser II. The V3-V4 region of the 16S rRNA gene was PCR amplified from 10 ng of genomic DNA using dual-indexed 16S rRNA Illumina iTags 341F (5′-CCTACGGGNBGCASCAG-3′) and 785R (5′-GACTACNVGGGTATCTAATCC-3′) supplemented with PNAs (Lundberg *et al*., 2012) designed to target host-derived amplicons from chloroplast and mitochondria 16S rRNA sequences (0.75 µM of each, PNABIO, Thousand Oaks, CA). Barcoded 16S rRNA amplicons were quantified using a Qubit dsDNA HS assay kit on a Qubit 3.0 fluorometer (Invitrogen, Carlsbad, CA), pooled in equimolar concentrations, purified using Agencourt AMPure XP magnetic beads (Beckman Coulter, Indianapolis, IN), quantified using a Qubit dsDNA HS assay kit on a Qubit 3.0 fluorometer (Invitrogen), and diluted to 10 nM in 30 μL total volume before being submitted to the QB3 Vincent J. Coates Genomics Sequencing Laboratory facility at the University of California, Berkeley for sequencing using Illumina Miseq 300 bp pair-end with v3 chemistry. V3-V4 16S rRNA amplicon sequencing reads were demultiplexed in QIIME2 (Bolyen *et al*., 2019) applying a minimum predicted accuracy of Q30. Chimera detection and removal were performed using DADA2 (Callahan *et al*., 2016) and high-quality amplicon sequence variants (ASVs) were assigned taxonomy using Silva database 138.2 SSU NR99.

### Time-course RNAseq

Plants inoculated with SRC2v4, HK, or with Mock, were harvested at 1 h, 24 h, 72 h, 1 week, 2 weeks, and 1 month post-inoculation. Harvests were performed around 12:30 PM for all time points within a 30-minute collection window of two boxes per treatment. Plants were placed in disinfected trays lined with new aluminum foil. Shoots and roots were separated using sterile scissors, and roots were gently cleaned with paper towels before immediate flash-freezing in liquid nitrogen. Shoots were also flash-frozen in liquid nitrogen. All tissues were stored at −80°C until nucleic acid extraction and sequencing. Total RNA was extracted from 10-50 mg tissue aliquots of bulk root or bulk shoot tissue from individual plants. Briefly, RNA was extracted via phase separation with TRIzol (Catalog No. 15596026; Invitrogen, Carlsbad, CA) followed by column purification. Resulting total RNA was treated to remove excess ribosomal RNA via polyA selection of mRNAs and sequenced on Novaseq S4 (Illumina) in a paired end 2x151nt run, yielding ∼30 million reads per sample. Paired-end reads were aligned to the Sorghum bicolor RTx430 v2.1 genome (Varoquaux *et al*., 2019) using the STAR aligner version 2.7.9a (Dobin *et al*., 2013). Read counts per gene were analyzed in edgeR and differential gene expression (DE) was tested using the quasi-likelihood framework (Robinson *et al*., 2010; Chen *et al*., 2016).

### Metabolomics

To characterize the metabolomic profiles of sorghum plants inoculated with SRC2v4, exometabolites were collected from the root and rhizosphere compartments. Experiments followed the general framework described above, with identical growth and sampling conditions. Harvests were performed one month post-inoculation, around 12:30 PM, within a 30-minute collection window, with four replicate microboxes per treatment. Rhizosphere samples were collected by adding 8 mL of epiphyte removal buffer (0.75% KH₂PO₄, 0.95% K₂HPO₄, 0.1% Triton X-100 in ddH₂O; filter-sterilized at 0.2 μm) to each sample, followed by manual agitation. The resulting rhizosphere was centrifuged at 3,700 rpm for 10 minutes to pellet the soil fraction (Fonseca-García *et al*., 2024).

Metabolite extracts (MEs) were analyzed by liquid chromatography–electrospray ionization tandem mass spectrometry (LC–ESI–MS/MS) using a Thermo Vanquish Flex UHPLC system coupled to a Thermo Q Exactive Plus Orbitrap mass spectrometer. For reversed-phase C18 chromatography, 5–10 µL of each extract were injected and separated on a Thermo Hypersil GOLD column (2.1 × 150 mm, 3 µm particle size) maintained at 40°C with a flow rate of 400 µL min⁻¹. The mobile phases consisted of (A) water with 0.1% formic acid and (B) acetonitrile with 0.1% formic acid, starting at 90:10 (A:B). The gradient proceeded as follows: 0–2 min, 90% A; 2–11 min, 10% A; 11–12 min, 10% A; 12–12.5 min, 10% A with an increased flow rate of 500 µL min⁻¹; 12.5–13.5 min, 90% A at 500 µL min⁻¹; 13.5–14 min, 90% A at 500 µL min⁻¹; 14–14.5 min, 90% A while decreasing flow rate to 400 µL min⁻¹; and 14.5–16 min, 90% A.

Mass spectrometry was performed in both positive and negative ionization modes using higher-energy collision dissociation (HCD). The heated ESI source parameters were as follows: spray voltage, 3.7 kV (positive) or 3.0 kV (negative); capillary temperature, 350°C; S-lens RF level, 50 a.u.; and auxiliary gas heater temperature, 150°C. Full MS scans were acquired at a resolution of 120,000 FWHM (m/z 200) over an m/z range of 80–800, with an automatic gain control (AGC) target of 3 × 10⁶ ions and a maximum injection time of 20 ms. Data-dependent MS² (dd-MS²) spectra were acquired at 15,000 FWHM, with an AGC target of 1 × 10⁵ ions, maximum injection time of 100 ms, isolation width of 0.4 m/z, loop count of 12, and normalized collision energies of 20, 40, and 60 eV.

For the generated data, processing was performed using Thermo Compound Discoverer 3.3. Metabolite identification was carried out using both internal reference libraries and external databases, including MzCloud, GNPS, and MoNet, among others. For reversed-phase (RP) positive and negative ionization modes, spectra were aligned using an adaptive curve with a maximum retention time (RT) shift of 0.3 minutes and a 3 ppm mass tolerance. Peaks were selected based on a minimum intensity threshold of 1.5 × 10⁴ and a chromatographic signal-to-noise ratio of 3. Detected features were grouped using a 3 ppm mass tolerance and a 0.3 min RT tolerance, and subsequently filtered based on sample occurrence. Compound assignment was based on isotopic pattern, RT, MS1, and/or MS2 data. All identifications and integrated peaks were manually validated before export for statistical analysis. Compound identifications were categorized into four confidence levels: 1. Match with MS1, MS2, and RT; 2. Match with MS2 only; 3. Match with MS1 and RT; 4. Match primarily with MS1 and partially with MS2 (**Supplement SD4**). Totals of 3750 polar features and 15107 non polar features were predicted including positive and negative ion modes across different treatments.

Principal component analysis (PCA) of metabolite profiles was conducted using *stats::prcomp* (R Core Team, 2023) function in R, with data log-transformed prior to analysis. Principal component analysis was performed using *vegan* (Dixon, 2003). All additional metabolite statistical analyses, including data normalization, transformation, and metabolite enrichment, were carried out using MetaboAnalyst v6.0 (Pang *et al*., 2024). Unless otherwise specified, analyses followed MetaboAnalyst’s default pipeline, with data integrity checks, missing value imputation by k-nearest neighbors (KNN), with log-transformation, and false discovery rate (FDR) correction for multiple testing.

### Metatranscriptomics

Experiments followed the general framework described above, with identical growth and sampling conditions. Rhizosphere samples were collected 1 month post-inoculation from two replicate microboxes per treatment, each containing four individually sampled plants with physically separated root systems, yielding eight independent biological replicates per treatment. Rhizosphere material was collected by adding 8 mL of epiphyte removal buffer to each sample, followed by manual agitation. Samples were then centrifuged at 3,700 rpm for 10 minutes to pellet the soil fraction and flash-frozen in liquid nitrogen. Total RNA was extracted from the resulting pellet using RNeasy PowerSoil Total RNA Kit (Qiagen), incorporating a freshly prepared phenol/chloroform/isoamyl alcohol solution (25:24:1 v/v, pH 6.5-8.0) to enhance lysis and RNA recovery. RNA was treated with DNase using TURBO DNA-free kit (Thermo Fisher Scientific) before downstream steps. Library preparation and sequencing were performed by UC Berkeley QB3 Genomics (Berkeley, CA, RRID:SRC_022170). Total RNA quality was assessed on an Agilent 2100 Bioanalyzer. We then enriched for mRNA using the Lexogen RiboCop Meta probes, and assessed the mRNA quality on Bioanalyzer. Libraries were prepared using the KAPA RNA Hyper Prep kit (Roche KK8581). Truncated universal stub adapters were ligated to cDNA fragments, which were then extended via 12 cycles of PCR using unique dual indexing primers into full-length Illumina adapters. Library quality was checked on an AATI (now Agilent) Fragment Analyzer. Library molarity was measured via quantitative PCR with the KAPA Library Quantification Kit (Roche KK4824) on a BioRad CFX Connect thermal cycler. Libraries were then pooled by molarity and sequenced on an Illumina NovaSeq X, 25B flowcell for 2 x 150 cycles, targeting at least 25M reads per sample. Fastq files were generated and demultiplexed using Illumina BCL Convert v4 and default settings.

All 15 SRC2v4 isolate genomes were concatenated into a single reference and indexed with *Salmon* (Patro *et al*., 2017). To assess relative transcriptional activities of the SRC2v4 members, transcripts per million (TPM) were assessed with *Salmon quant*. Average gene expression across all genes for each genome was calculated for all isolates in SRC2v4-treated rhizospheres.

## Results

### Four design strategies for SynCom complexity reduction

To determine the extent to which Syncom complexity can be reduced without sacrificing the collective benefit to the host, we engineered four reduced-complexity variants (**Fig. 1, Supplement SD1**) of the parent SRC1 consortium (Fonseca-García *et al*., 2024). Out of the original 57 member community, 15 were selected for inclusion in each of four distinct communities based upon maximization of four different criteria (see Methods): phylogenetic diversity (SRC2v1), predicted genomic functional capacity of the community (SRC2v2), positive correlation of colonization with increased plant biomass (SRC2v3), and isolate colonization of sorghum roots (SRC2v4). These four SRC2 variants were assessed with microbiome and host phenotypic measurements to test for successful colonization and plant growth promotion (PGP) activity.

### SRC2v1-4 exhibit variable community stability and plant growth promotion

To characterize the reduced SynCom variants, we deployed 16S rRNA amplicon metagenomics to interrogate community stability and sorghum root colonization. Alpha-diversity analysis revealed a clear and consistent pattern: all variants except SRC2v4 lost diversity upon entering the rhizosphere or root (**Fig. 2a, S2**). SRC2v2 and SRC2v3 were filtered at both compartments, while SRC2v1 was stable through the rhizosphere but collapsed in diversity at the root. SRC2v4 alone maintained the diversity of its inoculum across both compartments. Such observed root microbiome filtration effects are well documented (Lundberg *et al*., 2012; Bulgarelli *et al*., 2012; Edwards *et al*., 2015; Xu *et al*., 2018) and provided the first indication of SRC2v4 robustness.

Exploration of the beta-diversity helped sharpen this picture. PCoA of all four SynComs resolved two distinct clusters: SRC2v1/SRC2v4 and SRC2v2/SRC2v3 which persisted in both the rhizosphere and root, compartments directly under host influence (**Fig. S3**). Strikingly, these differences dissolved entirely in bulk soil (PERMANOVA R²=0.25, p=0.16). Together, these results demonstrated that community differentiation was not intrinsic to the inocula themselves, but emerged from the selective pressures of root exudates and physical colonization of the root interior.

An examination of the membership-level data across treatments helped to explain the observed filtration and differentiation effects (**Fig. 2b**). We detected a small number of ASVs in the SRC2 variant treatments that did not correspond to any inoculated strain (grey segments at the base of each bar, labeled “Other”). In total, this category comprised 20 bacterial ASVs spanning 13 genera (**Fig. S4**). The signal was dominated by four *Pantoea* and *Pseudomonas* ASVs, which together accounted for ∼90% of “Other” read counts and were detected across all treatments, including the Mock. Given the well-documented prevalence of *Pantoea* – and to a lesser extent *Pseudomonas* – in seed microbiomes, and the higher abundance of these ASVs in the endosphere relative to soil, we infer that they represent seed endophytes that escaped sterilization. The remaining non-target taxa occurred at low abundance and included a single *Streptomyces* ASV detected across all four variants (∼8% of “Other” reads; absent from the Mock), consistent with a potential media contaminant.

Among all of the SynCom variants, *Bacillus* isolates were consistently poor colonizers while *Pseudomonas* grew robustly in SRC2v2 and SRC2v3 but were suppressed in SRC2v1 and SRC2v4 suggesting community-dependent colonization. By contrast, isolates selected for their capacity to metabolize the sorghum root exudates (Yang *et al*., 2004; Weston *et al*., 2013; Fonseca-García *et al*., 2024) were dominant root colonizers: strikingly, SO8-2, which was abundant in the root and rhizosphere across all four SynComs, along with SO6b and PH2, accounted for 46% of the SRC2v1 and SRC2v4 communities *in planta*. Taken together, the divergent responses of the above-listed taxa drive beta-diversity differences observed between SRC2v2/SRC2v3 and SRC2v1/SRC2v4 clusters (**Fig. S3**) and illustrate the filtration effects of sorghum roots on SynCom community cohesion.

Differential abundance testing reinforced SRC2v4’s exceptional stability (**Fig. S5**). Relative to inoculum composition, SRC2v1 enriched 5 members *in planta*, SRC2v2 enriched 9, and SRC2v3 shifted at least 2. SRC2v4 shifted none. Across all variants, SRC2v4 was the most compositionally representative of its inoculum, a crucial trait for reproducible, controlled microbiome treatments.

Finally, we sought to identify the impact of each SynCom variant on plant host performance. Germinating seedlings were inoculated with each one of the four variants or a Mock; they were then grown for one month in gnotobiotic conditions, then harvested and phenotyped. SRC2v1 failed to elicit significant PGP in any phenotype; by contrast, SRC2v2, SRC2v3, and SRC2v4 significantly increased shoot fresh weight, dry weight, shoot length, and water content (**Fig. 3**), albeit to variable magnitudes. While no statistically significant differences were detected among SRC2v2, SRC2v3, and SRC2v4 for the majority of traits, SRC2v4 consistently yielded the highest median scores for three of the four measured parameters. Collectively, these data demonstrated that SynCom variants of equal size (n=15) and treatment biomass (∼1g) do not always induce similar phenotypic outcomes in the host and that SRC2 membership determines PGP outcomes in the host.

To select a single SynCom for further molecular characterization, we considered the colonization and PGP characteristics of all SRC2 variants. In terms of root and rhizosphere colonization, all variants exhibited diversity reduction in roots relative to inoculum. However, in the rhizosphere, only SRC2v1 and SRC2v4 retained diversity equal to the starting inoculum (**Fig. S2**) and greater diversity than both SRC2v2 and SRC2v3 (**Fig. 2a**). In terms of plant phenotypes, SRC2v4 exhibited greater (although not statistically significant) median values for shoot length, shoot fresh weight, and shoot dry weight relative to SRC2v1, and SRC2v4 significantly improved shoot water content relative to SRC2v1 (**Fig. 3**). Considering all these results together, we selected the top performing variant for further study: SRC2v4.

### Metatranscriptomics identifies active SRC2v4 members in the rhizosphere

To identify the most active members of the SRC2v4 SynCom in sorghum rhizospheres, we quantified gene expression using a metatranscriptomics approach. The relative transcriptional activity of each isolate was normalized to the number of genes in each genome (**Fig. S6**). These data revealed that several members of the community showed almost no transcription activity (SAI104, TBS030, SO6a), while the top 7 most transcriptionally active members accounted for over 90% of the total TPM/gene in the community. Additionally, we observed that the most dominant member of the community in terms of activity, *Pseudomonas* TBS028, was only the 5th most abundant member of the community as assessed by 16S rRNA (comprising just 9.9% of total 16S reads in the rhizosphere, data derived from experiments shown in (**Fig. 2, S8, Supplement SD5**). Despite this observation, we also observed that mean transcriptional activity significantly correlated with abundance (Spearman’s ρ=0.70, p=0.005). Collectively, transcriptional analysis of rhizosphere activity revealed a community where abundance tended to explain activity that was largely dominated by the top 50% most active isolates.

### Live SRC2v4 inoculation is required for plant growth promotion

While SRC2v4 proved effective in terms of PGP, we did not yet know if this could be attributed to sustained interactions between the plant and living microbes. For instance, the mere presence of Microbe-Associated Molecular Patterns (MAMPs) like flagellins, peptidoglycans, and lipopolysaccharides recognizable to the plant immune system (Boller & Felix, 2009; Macho & Zipfel, 2014) might be sufficient to induce the observed PGP. To rule out the latter possibility, we compared sorghum phenotypes upon inoculation with either SRC2v4, HK, or Mock (**Fig. 4a**). Live SRC2v4 treatment increased shoot fresh weight, shoot dry weight, and root fresh weight. Conversely, HK treatment failed to significantly induce any PGP, despite exhibiting phenotypes intermediate to Mock and SRC2v4. Considering these observations, we conclude that while application of HK particles induced minor effects, host interaction with live SRC2v4 was required for PGP.

As the SynCom variants were each derived from the original SRC1 community, we next sought to compare the phenotypic impact of SRC2v4 relative to SRC1. We again measured above and below ground phenotypes one month after inoculation with either SRC2v4, SRC1, or Mock treatments under gnotobiotic conditions (**Fig. 4b**). Both SRC1 and SRC2v4 increased shoot fresh weight, dry weight and shoot length compared to the Mock. Importantly, SRC2v4 failed to match PGP mediated by SRC1, indicating that our derivative community with reduced complexity retained beneficial effects of the original SRC1, but with reduced magnitude.

### Sorghum’s dynamic, spatiotemporal transcriptional response to SRC2v4

To mechanistically dissect SRC2v4’s impact, we profiled the host transcriptional response with a one-month, six-timepoint RNA-seq time course including SRC2v4, HK, and Mock treatments. SRC2v4 induced significant differential expression (DE) in 613 unique genes compared to Mock across all tissues and timepoints (**Fig. 5a**). This response displayed a complex spatiotemporal dynamic: strong initial DE localized to roots at 1 day, followed by a systemic wave of DE in shoots by 3 days, frequently involving the same genes (**Fig. 5b**). Finally, at the 1 month timepoint, the dominant DE signal resided in the root. In total, 73 genes exhibited a signature of early root regulation coupled with subsequent shoot DE. This time-resolved transcriptional pattern demonstrates a complex host accommodation to the live SRC2v4 community.

Live SRC2v4 treatment and HK also induced divergent host transcriptional responses. The HK treatment elicited a transient DE pulse, peaking in the root at 1 day (**Fig. S7**), but failed to induce significant DE in the shoot at any point or or sustained DE in the root. Conversely, SRC2v4 established a dynamic, sustained spatiotemporal DE pattern: strong induction began in the root at 1 day, translated systemically to the shoot at 1 week, and returned to a dominant DE signal in the root at 1 month. This demonstrated that while dead microbial components triggered an early, transient root response, the live, colonizing SRC2v4 community amplified and sustained the root response relative to HK and led to additional transcriptional reprogramming in the shoot.

To resolve if the root-to-shoot transcriptional shift induced by SRC2v4 was mediated by signaling or reflected delayed microbial localization of the shoot, we employed 16S rRNA amplicon metagenomic profiling. Due to plant tissue mass limitations at time of collection, sample aliquots for 16S rRNA in addition to the RNAseq aliquots could only be collected at 1 week, 2 week, and 1 month timepoints. Across the sampled dataset, SRC2v4-associated ASVs were among the most highly observed and were found in each compartment and treatment group (**Fig. S8a-b**), demonstrating that the inoculated isolates were detectable above environmental and low-abundance sequencing noise. The live SRC2v4 treatment established stable and substantial colonization in both root and shoot tissues from 1 week through 1 month post-inoculation (**Fig. S8c, S9a**); by contrast, HK and Mock yielded relatively low detection of SRC2v4 members (consistent with residual dead DNA or contamination, respectively), and dominance of ASVs in the “Other” category, representing both putative seed endophytes and organelle genome-derived reads (**Fig. S9b**). Combining these data with the host transcriptomics analysis (**Fig. 5**), we conclude that the observed shift in host DE in shoots is likely directly coupled to systemic colonization of the live SRC2v4 community, rather than root-mediated transcriptional signaling.

To further characterize the dynamic, spatiotemporal DE induced by SRC2v4, we assessed the 73 genes exhibiting a root-to-shoot DE signature, revealing a core functional response driven by hormone regulation and secondary metabolism. *Brassinosteroid Insensitive 1-Associated Receptor Kinase 1* (BAK1), a key regulator of brassinosteroid signaling and microbial perception (Li & Chory, 1997; He *et al*., 2000; Li *et al*., 2002; Nam & Li, 2002)), displayed the strongest overall DE (log_2_FC=11.8, shoot, 1 week) and a representative spatiotemporal profile (**Fig. 6a**). Furthermore, three distinct genes known to modulate availability of the plant growth hormones indole acetic acid (IAA) and cytokinin (CK) were upregulated in roots at day 1 including: *Cytokinin-O-glucosyltransferase* (Pineda Rodo *et al*., 2008), *Cytokinin-N-glucosyltransferase* (Šmehilová *et al*., 2016), and *Methyl Esterase 1* (Yang *et al*., 2008) (**Fig. S10**). Additionally, functional enrichment analysis confirmed that the flavonoid biosynthetic pathway was strongly regulated, marked by the enrichment of *Chalcone Synthase* (CHS) (**Fig. 5c**) and the concerted upregulation of downstream enzymes: *Chalcone Isomerase* (CHI), *Flavonol Synthase* (FLS), and *Dihydroflavonol 4-Reductase* (DFR) (**Fig. 6b**). Collectively, the transcriptional data define dynamic, spatiotemporal plant-microbe interactions with a strong emphasis on host-mediated flavonoid production and hormone regulation.

### Metabolomics confirms enrichment of flavonoids in SRC2v4-treated roots

To validate the flavonoid-centric host transcriptional signature, we characterized the root and rhizosphere metabolomes with LC–ESI–MS/MS one month after inoculation. While Mock and SRC2v4 treatments were significantly differentiated in both compartments, the treatment effect was more pronounced in the roots (**Fig. 7a**; PERMANOVA: root R^2^=0.22, p=0.001; rhizosphere R^2^=0.16, p=0.001). In the rhizosphere, SRC2v4 treatment enriched 38 metabolites and depleted 15 (**Fig. 7b**), a shift defined by the accumulation of signaling lipids and flavonoids. The most highly enriched compound was Flavone + 1O; OMe (Log_2_FC=5.16). Various enriched lysophosphatidylethanolamines (LPEs 19:1, 17:1, 16:1, and 16:0) suggested active membrane remodeling or lipid-mediated signaling. The more robust root response featured 211 enriched and 94 depleted metabolites (**Fig. 7b**). Notably, five enriched flavones in the root in addition to Flavone + 1O; OMe in the rhizosphere (**Fig. 7c**) support the observed upregulation of flavonoid biosynthetic transcripts. Interestingly, depletion of Hydroxycoumarin + O-Hex, which shares phenylpropanoid precursors with flavones, suggested that the roots actively tune carbon flux across these pathway branches. The persistence and accumulation of these metabolites one month after the peak gene upregulation (3 days, **Fig. 6b**) demonstrates a coordinated, sustained redirection of carbon and lipid resources toward specialized metabolites to facilitate long-term SRC2v4 colonization.

Finally, correlation analysis integrating metabolite profiles with 16S rRNA amplicon metagenomic data revealed associations between microbial composition and metabolite signatures (**Fig. S11**). Putative sorghum endophytes (“Other”) showed significant negative correlations with 30 enriched and four depleted metabolites, a pattern also observed for SO6a. In contrast, SRC2v4 members SO6b and PH2 positively correlated with several flavone-related metabolites. These patterns are consistent with SRC2v4-mediated restructuring of the root microbiome and metabolome, where enrichment of SRC2v4 members coincides with flavonoid-associated metabolite shifts and reduced abundance of native endophytes. However, when the analysis was restricted to SRC2v4-inoculated samples, no significant associations were detected, indicating that metabolite variation was primarily driven by the presence of the SRC2v4 community rather than differences in the abundances of individual strains. Collectively, these results suggest that SRC2v4 inoculation reshapes the root metabolome at the community level and is associated with persistent enrichment of flavonoid-related metabolites.

## Discussion

### Design strategies

In this study, we produced a reduced-complexity, yet beneficial SynCom derived from our previously published 57 bacterial member SynCom, SRC1. We utilized distinct design strategies in an attempt to test their abilities to recapitulate the phenotypic benefits to the sorghum host that were observed in SRC1 treatment. Our SynCom variant isolate selections designed to maximize predicted functional capacity (SRC2v2), abundance correlation with desired phenotypes (SRC2v3), and host colonization (SRC2v4) all improved PGP as observed in other systems (Zhuang *et al*., 2021; Afrizal *et al*., 2022; Berihu *et al*., 2023). While there were minor differences in median plant growth promotion across SRC2v2, SRC2v3, and SRC2v4, none of these were significant. The failure of SRC2v1 – a variant selected without reference to host interaction data – is an instructive negative result. Despite maintaining rhizosphere diversity comparable to SRC2v4, SRC2v1 collapsed in the root and elicited no significant PGP, demonstrating that phylogenetic breadth alone is insufficient as a design criterion. That the three variants incorporating host-derived selection data (colonization capacity, exudate utilization, or phenotypic association) all succeeded where the taxonomically-motivated variant failed argues that functional compatibility with the host, rather than ecological representativeness, is the primary determinant of SynCom efficacy. Finally, we note that the high-complexity, 57 member SRC1 community outperforms the top-performing reduced-complexity variant, SRC2v4 (**Fig. 4**). This result suggests the observed reduction in performance was due to either the greater size and complexity of SRC1 or the omission of key microbes (van der Heijden *et al*., 2006; Banerjee *et al*., 2018) when sub-selecting the SRC2v4 variant. Indeed, single isolate or genus SynComs have been found solely responsible for host impacts in complex SynComs (Niu *et al*., 2017; Kwak *et al*., 2018; Finkel *et al*., 2020). Ultimately, these results highlight the efficacy of multiple SynCom design strategies, the effect of taxonomic complexity on PGP benefits, and the general value of SynCom application.

### Living inocula establish sustained commensal relationships with host sorghum

Our dense, longitudinal transcriptional sampling of sorghum response to SynCom application enabled deep characterization of microbiome establishment to its plant host. First, we wanted to differentiate the response to live SRC2v4 and HK. Consistent with typical transcriptional responses to purified MAMPs (Denoux *et al*., 2008; Stringlis *et al*., 2018), we observed a strong, early transcriptional response to HK in the root (**Fig. S7**). Likewise, SRC2v4 live inoculum induced an early peak in transcriptional activity. While beneficial bacterial mono-associations often trigger rapid responses that fade within 6 hours (Stringlis *et al*., 2018), our observation of peak DE between 1 and 3 days (**Fig. 5a**) aligns closely with commensal colonization dynamics (Tzipilevich *et al*., 2021). The early and dynamic responses to HK and SRC2v4 treatments mirrored plant responses to commensal *Pseudomonas* WCS417 and its flg22 peptide (Stringlis *et al*., 2018), suggesting an initial conserved recognition phase. The pulsed early activation and late reactivation we observed in roots points toward a more complex regulatory feedback loop. This complexity extends systemically to shoots. Similar to the DE overlap between root and shoot observed by (Montoya *et al*., 2025), we identified shared DE genes between the root and shoot at 3 days (**Fig. 5b**). Ultimately, these findings suggest that while early root transcription follows MAMP-like recognition, the plant eventually transitions into a stable, colonized state characterized by long-term transcriptional signatures (at least 1 month long) consistent with established SynCom-treated root models (Teixeira *et al*., 2021) effectively balancing acute perception with a sustained commensal equilibrium.

### Host transcription of BAK1 reflects complex growth and defense signaling

The sorghum transcriptome response to SRC2v4 follows a dynamic, spatiotemporal pattern. This general pattern was exemplified by BAK1, the most robustly regulated gene in our analysis (log_2_FC=11.8, shoot, 1 week post-inoculation; **Fig. 7a**). Consistent with previous observations of SynCom-induced priming (Paasch *et al*., 2023), BAK1 expression followed a distinct pattern: an initial pulse in the roots within 48 hours, followed by a sustained activation in the shoot from 3 to 7 days, and a terminal induction in the roots at 1 month. As a promiscuous co-receptor for both the pattern recognition receptor FLS2 (Heese *et al*., 2007; Chinchilla *et al*., 2007) and the brassinosteroid receptor BRI1 (Li & Chory, 1997; He *et al*., 2000; Li *et al*., 2002; Nam & Li, 2002), BAK1 sits at the critical nexus of the growth-defense tradeoff (Wu *et al*., 2025). The magnitude and duration of its upregulation suggests that SRC2v4 may be more than a passive occupant of the endosphere; rather, it may actively deploy molecular mimics or effectors to “tune” BAK1 activity. In addition, the enrichment of LPE metabolites in both root and rhizosphere could play a role in this signaling as they have been found to collaborate with brassinosteroids to stimulate root growth (Jeong *et al*., 2012). By inducing transcription of this co-receptor, the community could potentially promote growth-sustaining brassinosteroid signaling while simultaneously desensitizing the PRR complex – perhaps via the sequestration of BAK1 away from defense-active complexes – to circumvent an inhibitory immune response. This dual-purpose manipulation of BAK1 throughout our longitudinal observations provides one putative mechanistic basis for the observed PGP response to SRC2v4 application.

### A role for flavonoids in SRC2v4 plant-microbe interactions

The observed significant up-regulation of genes within the core flavonoid biosynthetic pathway (including those downstream of CHS CHI, and FLS) in SRC2v4-treated roots highlights a critical mechanism of host control, though its function likely differs fundamentally from its role in canonical legume-Rhizobia symbiosis. In legumes such as *Medicago truncatula*, the local exudation of early flavonoids (chalcones and flavanones) operates as a high-fidelity, reciprocal signaling mechanism: the host sends a species-specific molecular key (the flavonoid) that unlocks the bacterial *Nod* genes, leading to the production of Nod factors and subsequent nodule formation (Peters *et al*., 1986; Redmond *et al*., 1986; Shumilina *et al*., 2023). This interaction is primarily one of induction and specific recruitment. Our non-legume host sorghum plants do not possess the capacity for nodule organogenesis, suggesting that the observed increase in flavonoid synthesis serves a different ecological purpose than inducing nodulation. Notably, a recruitment role for flavonoids has been documented outside of the legume-nodulation interaction (Yu *et al*., 2021; He *et al*., 2022). Late-stage flavonoids including flavones, flavonols, anthocyanins, dihydroflavonols, and their derivatives, known for their broad antimicrobial and defense properties, are secreted into the rhizosphere and root tissues to establish a selective chemical barrier (Yan *et al*., 2022; Kumar *et al*., 2023). It is possible that strongly colonizing SRC2v4 members possess tolerance or detoxification mechanisms against these compounds. Roles for both recruitment and prevention of colonization by plant-produced flavonoids remain open. Future work will be required to distinguish between, or validate concurrent roles for these compounds in the SRC2v4 sorghum system.

Interestingly, the original, 57 member SRC1 elicited downregulation of lignin biosynthesis, a pathway linked to flavonoid biosynthesis (Fonseca-García *et al*., 2024). While the transcriptional signature of lignin precursor pathway downregulation in SRC1 was not recapitulated in SRC2v4 via transcriptomics or metabolomics, enrichment of flavonoids connects these two induced phenotypes in the host. Flavonoid and lignin precursor biosynthetic pathways both begin with the upper phenylpropanoid biosynthesis pathway. CHS uses *p*-coumaroyl-CoA and three malonyl-CoA to produce chalcone. It serves as the branchpoint and first committed step from monolignol biosynthesis toward flavonoids. As such, increasing CHS activity diverts carbon flux from the phenylpropanoid pathway, decreasing production of monolignol lignin precursors and increasing flavonoid production (Mahon *et al*., 2022). Increasing regulatory gene expression for flavonoid biosynthesis tends not only to increase flavonoids, but also deplete lignin (Shi *et al*., 2022). Evidence of such pathway tuning was observed in the depletion of Hydroxycoumarin + O-Hex, which shares phenylpropanoid precursors with flavones, further supporting the notion of host-mediated carbon flux modulation across these branches. Given the interconnectedness of these pathways and their regulation, it is possible that the lignin decreases upon SRC1 treatment and the increased flavonoid production by SRC2v4 treatment may share a common mechanism induced by shared bacteria in both SynComs.

## Conclusions

Rational reduction of the 57-member SRC1 consortium yielded SRC2v4, a stable, 15-member SynCom that preserves plant growth promotion in sorghum and reveals previously undescribed dimensions of host-microbe dialogue. The selection strategy of maximizing root colonization capacity proved superior to phylogenetic, functional genomic, or phenotype-correlation approaches, demonstrating that *in planta* persistence is a primary determinant of functional SynCom performance. Colonization by living SRC2v4 members – but not heat-killed inocula – triggered a spatiotemporally coordinated transcriptional response spanning root and shoot, anchored by the growth-defense hub BAK1 and culminating in sustained flavonoid biosynthesis. Corroborating metabolomics confirmed the durable accumulation of flavones in colonized roots, whose positive correlation with dominant *Flavobacteria* and *Enterobacter* members points toward a flavonoid-mediated microbiome structuring mechanism operating independently of canonical legume-Rhizobia signaling. Together, these findings establish a replicable framework for designing minimal, soil-independent microbial biostimulants and identify the host flavonoid pathway as a tractable molecular target for engineering more effective, next-generation plant-microbiome products for sustainable agriculture.

## Acknowledgements

This work was supported by US Department of Agriculture (CRIS 2030-12210-003-000D), USDA-NIFA (2019-67019-29306), and is a contribution of the Pacific Northwest National Laboratory (PNNL) Secure Biosystems Design Science Focus Area “Persistence Control of Engineered Functions in Complex Soil Microbiomes” (operated by the U.S. DOE under contract DE-AC05-76RL01830). A portion of this work was performed at the William R. Wiley Environmental Molecular Sciences Laboratory (EMSL), a national scientific user facility sponsored by the BER and located at PNNL. PNNL is a multi-program national laboratory operated by Battelle for the DOE under Contract DE-AC05-76RL01830.

## Author Contributions

DP: conceptualization, data curation, formal analysis, investigation, methodology, visualization, writing – original draft, review, and editing;. CF-G: conceptualization, data curation, formal analysis, investigation, methodology, visualization, writing – original draft, review, and editing; GK: methodology, formal analysis; HP: formal analysis, writing – review and editing; TW: conceptualization, funding acquisition, methodology, writing – review and editing; CC: methodology, formal analysis; PH: conceptualization, funding acquisition, investigation, supervision, writing – review, and editing; RE: conceptualization, funding acquisition, investigation, supervision, writing – review, and editing; DC-D: conceptualization, formal analysis, funding acquisition, investigation, supervision, writing – original draft, review, and editing.

## Data Availability

All genomes, 16S rRNA amplicon sequencing, plant RNAseq, and rhizosphere metatranscriptomics raw data files can be accessed through NCBI BioProject PRJNA1404338. Liquid chromatography-mass spectrometry (LC-MS) raw data files are openly accessible at the Mass Spectrometry Interactive Virtual Environment (MassIVE) community repository (http://massive.ucsd.edu ) under the accession MSV000100573. All scripts used are accessible on GitHub (https://github.com/deanpettinga/src2_paper).

## Supplemental Figures

**Figure S1.**
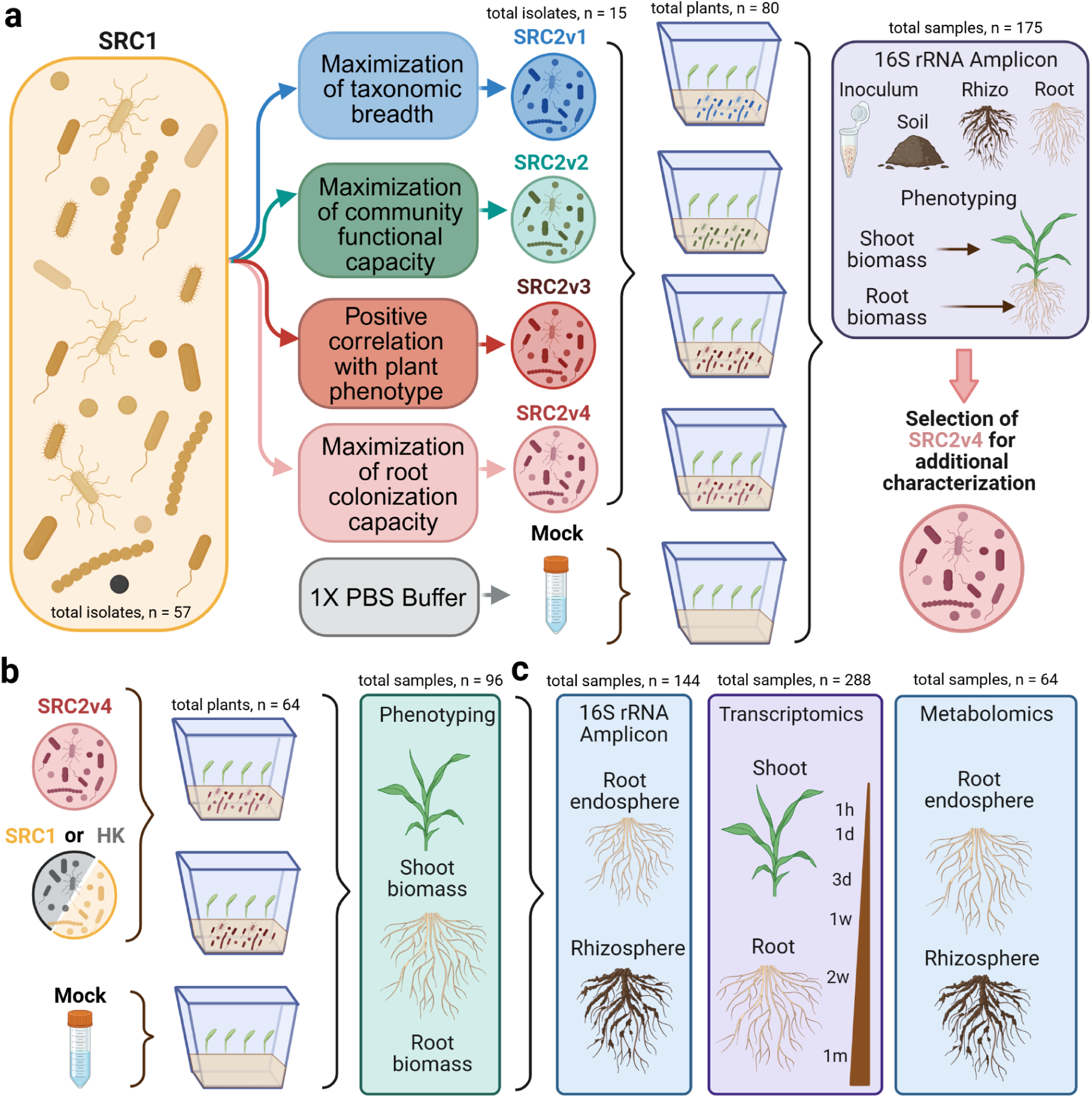
Experimental design of SRC2 SynCom construction and evaluation. **a)** The original sorghum consortium SRC1 (57 bacterial members) was reduced to 15-member SynCom using four complementary design strategies. SRC2v1 maximized taxonomic breadth by selecting isolates representing the phylogenetic diversity of SRC1. SRC2v2 maximized predicted functional capacity by selecting genomes that collectively retained the largest number of orthogroups across SRC1 genomes. SRC2v3 was formulated using results from a permutational membership experiment in which the most abundant genera in SRC1 (*Bacillus*, *Pseudomonas*, and *Streptomyces*) were systematically depleted, and strains showing the strongest positive associations between abundance and plant shoot biomass were prioritized. SRC2v4 was designed to maximize root colonization capacity by selecting the 15 strains with the highest cumulative relative abundance across root samples. Each SRC2 formulation was inoculated onto Sorghum bicolor (cv. RTx430) seedlings grown in sterile microbox assays, alongside mock buffer controls. Plants were grown under gnotobiotic conditions for one month and evaluated for plant phenotypes and microbial colonization. Shoot and root biomass measurements were collected, and microbial community composition was assessed using 16S rRNA amplicon sequencing of inoculum, soil, rhizosphere, and root samples. Based on these results, SRC2v4 was selected for further characterization. **b)** Follow-up experiments with SRC2v4 included comparisons with heat-killed inoculum (HK) and the original SRC1 consortium. Plants were grown under gnotobiotic conditions for one month and evaluated for above and below-ground plant phenotypes. **c)** SRC2v4 functional analyses were performed to assess microbial transcriptional activity and host responses. These experiments included time-course host transcriptomics (1 h to 1 month post-inoculation), microbial community profiling of roots and shoots using 16S rRNA amplicon sequencing, root and rhizosphere metabolomics, and rhizosphere metatranscriptomics. Plants were grown under the same gnotobiotic conditions and harvested up to one month post-inoculation.

**Figure S2.**
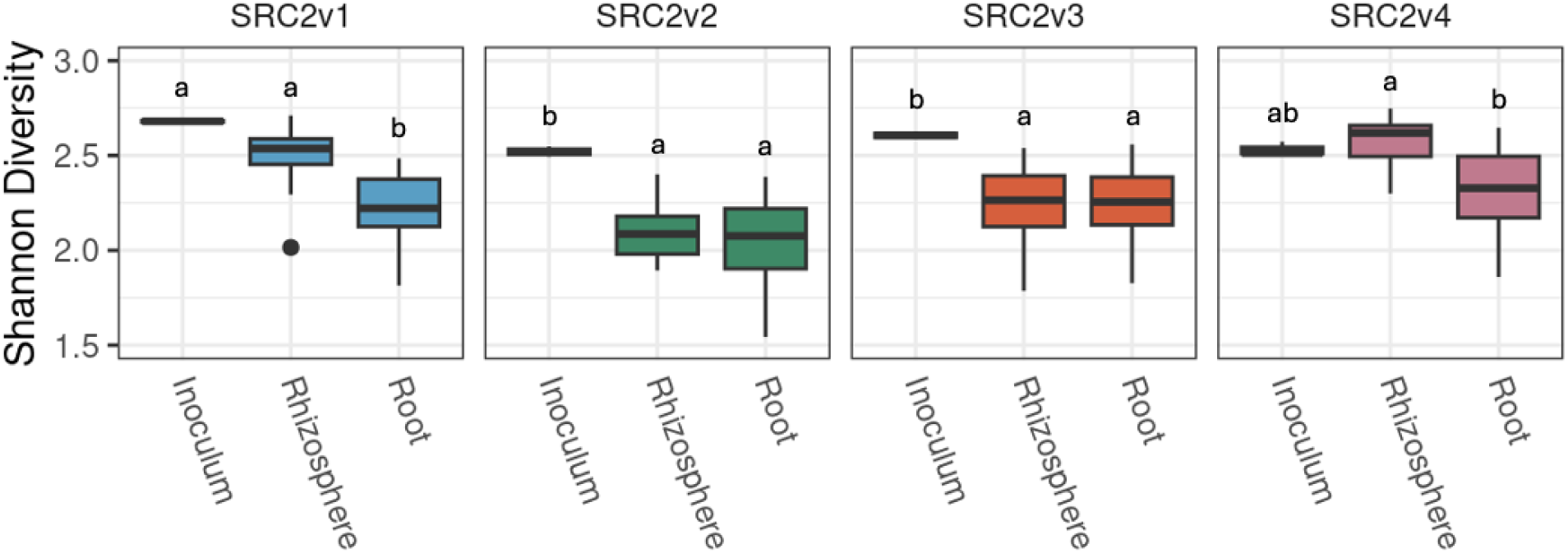
Shannon Index across colonization compartments illustrates variable filtering effects of host plant on diversity between SynCom treatments. Stability assessment of each SynCom across colonization compartments demonstrates the degree of host filtering by tracking diversity differences comparing the inoculum to rhizosphere and root. All communities showed a significant loss of diversity from the inoculum to the root confirming host filtering, but SRC2v4 was robust to diversity loss from inoculum to rhizosphere before exhibiting some filtering from rhizosphere to root. In all panels, letters above the boxplots represent statistical significance based on one-way ANOVA followed by post-hoc Tukey’s HSD (alpha = 0.05).

**Figure S3.**
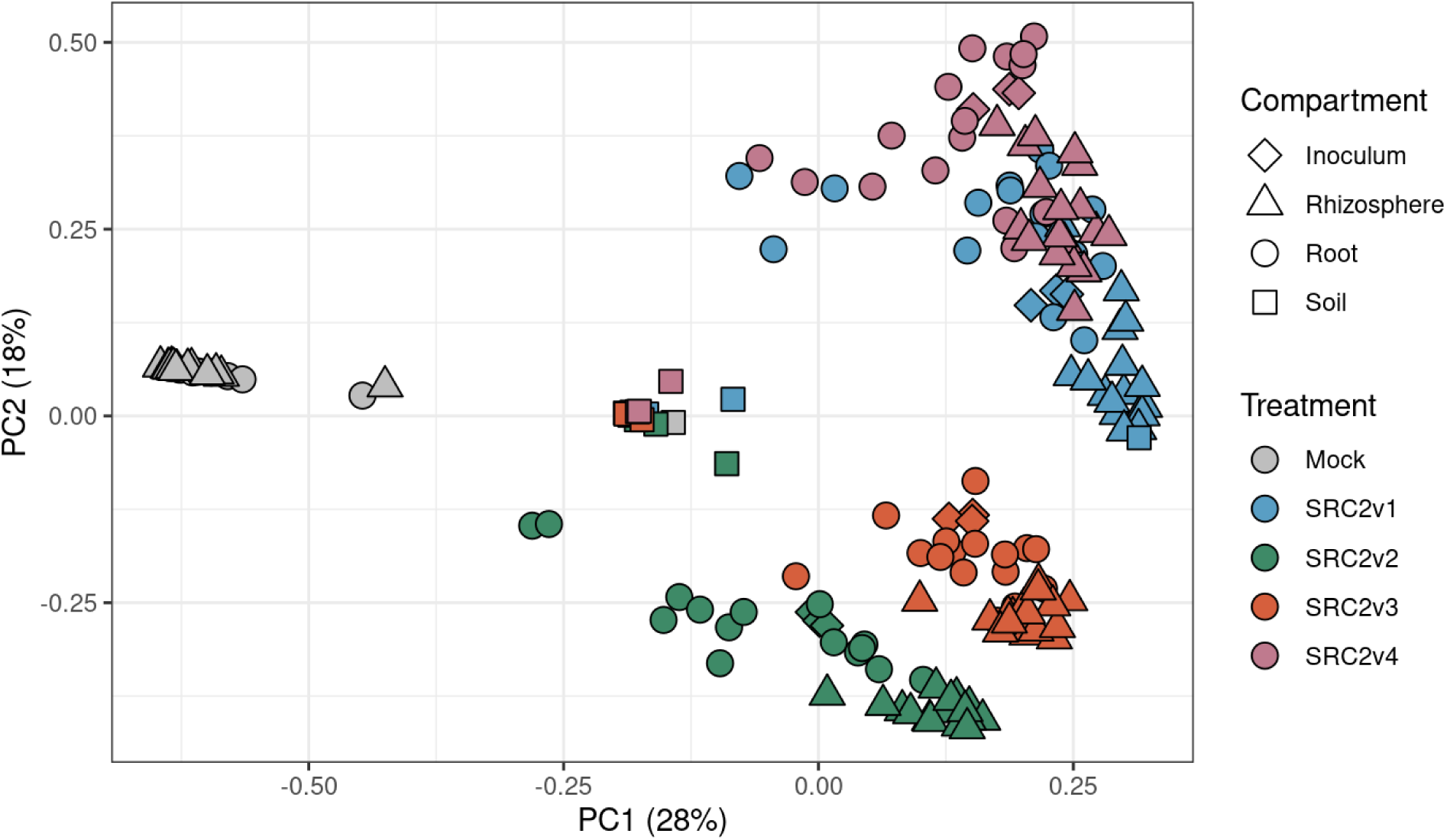
SynComs SRC2v1-4 colonize sorghum and elicit variable host growth phenotypes. SRC2v1-4 exhibit variable increases to plant host phenotypes and colonization. Surface-sterilized, germinating seedlings were inoculated with each SRC2 formulation, planted in sterilized 5L microboxes with 1kg of calcined clay. After 1 month, samples were harvested for 16S amplicon metagenomics and host phenotyping. Beta-diversity of SRC2v1-4 colonization in root, rhizosphere and bulk soil visualized by PCoA reveals two clusters: SRC2v1+SRC2v4 and SRC2v2+SRC2v3.

**Figure S4.**
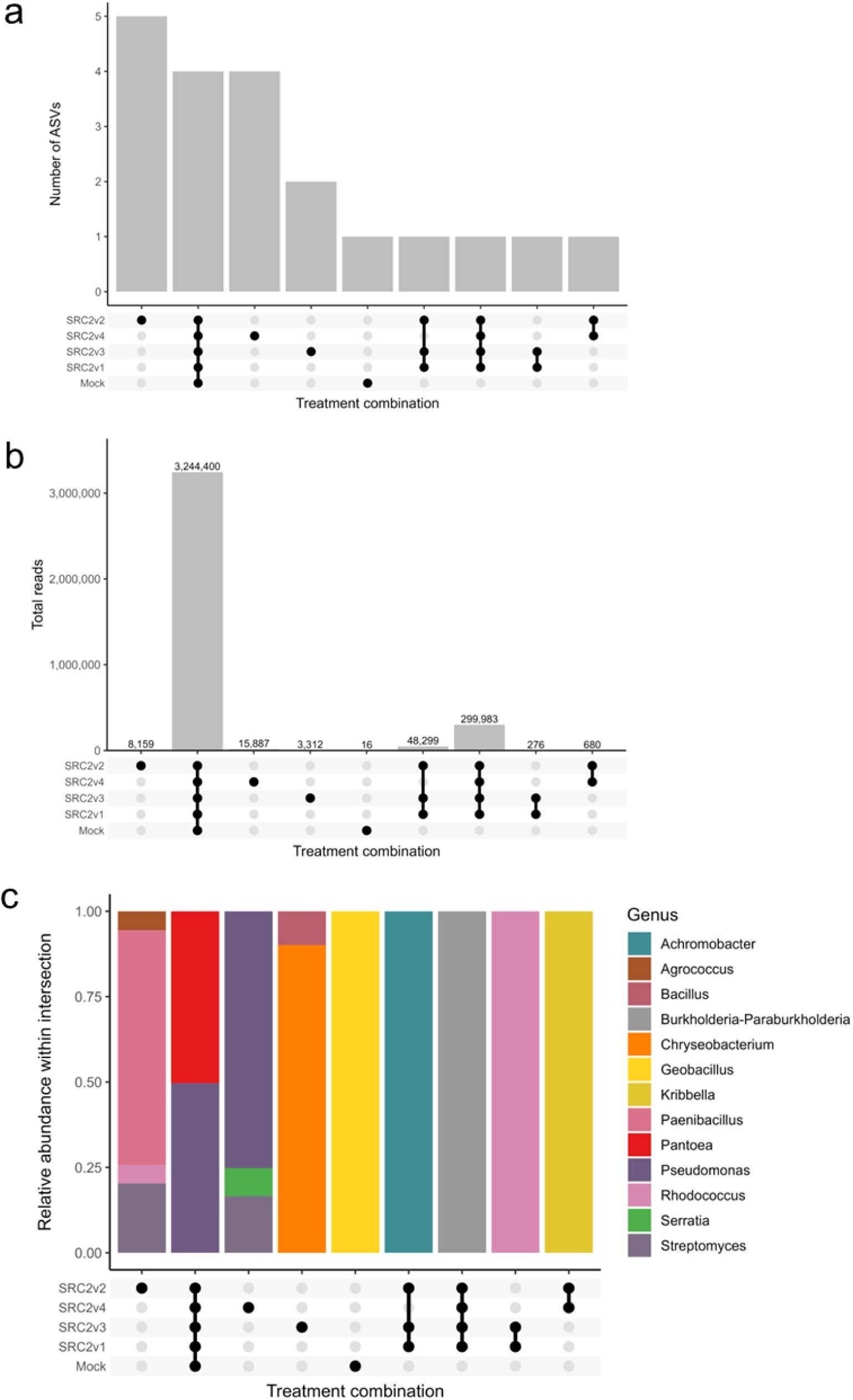
Distribution and taxonomic composition of all ASVs not matching intentionally included strains from the SRC2 variants detected across the experiment in any sample type. Only ASVs with ≥100 total reads and detected in ≥4 replicates within a treatment were retained for this analysis. **a)** UpSet plot showing the number of non-SynCom derived ASVs shared among treatments (Mock, SRC2v1-SRC2v4) across plant compartments (root, rhizosphere, and inoculum). **b)** Total read counts contributed by the ASVs shown in a) for each treatment combination across root, rhizosphere, and inoculum samples. Numbers above bars indicate the summed reads per intersection. **c)** Relative abundance of the taxonomic composition of the ASVs included in the intersections in a) at genus level.

**Figure S5.**
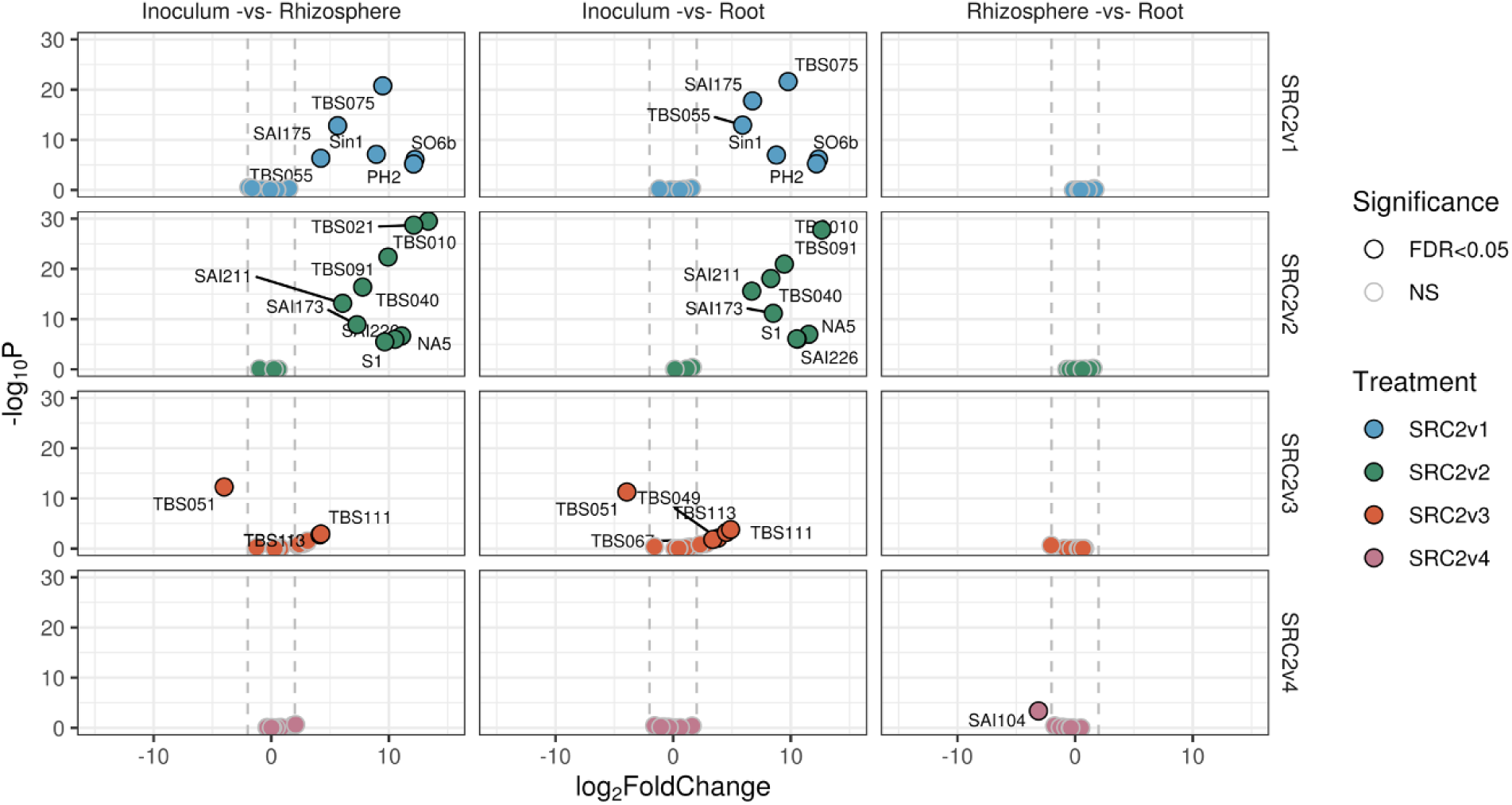
SRC2v1-4 exhibit variable member abundance stability from inoculum to rhizosphere and root. Volcano plots comparing Rhizosphere-Inoculum, Root-Inoculum, and Root-Rhizosphere identify members of each SynCom with differential abundance between compartments. All SynComs are stable between Root and Rhizosphere. In contrast to the starting Inoculum, about half of the members of SRC2v1 and SRC2v2 increase in abundance when colonizing plant compartments. Only two members SRC2v3 vary in relative abundance between inoculum and plant compartments whereas SRC2v4 remains stable.

**Figure S6.**
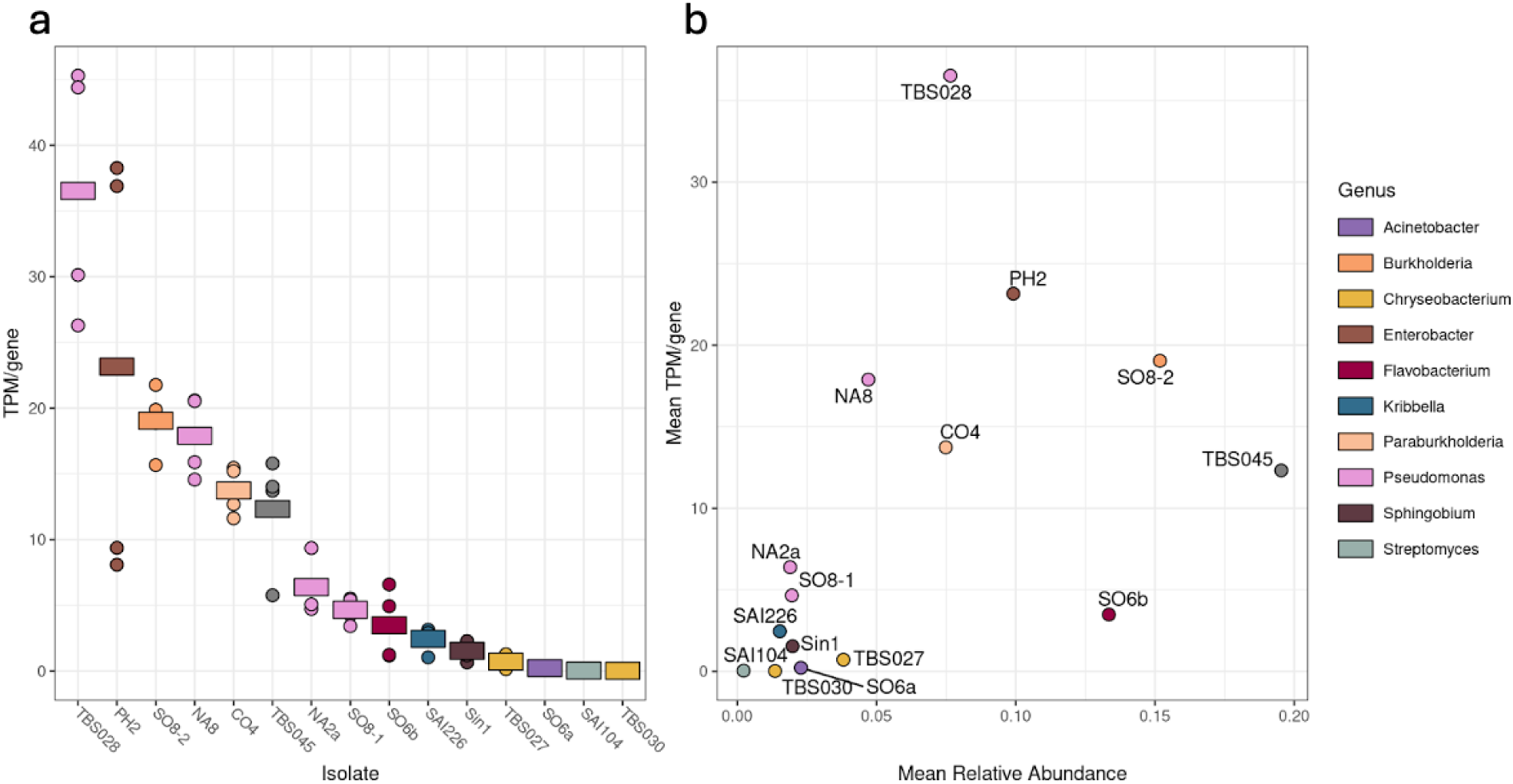
Metatranscriptomic profiling of SRC2v4-treated sorghum rhizospheres reveals variable transcriptional activity among isolates. For each isolate, gene transcription was quantified by *Salmon*. **a)** Average gene expression across all genes in each genome was calculated for all isolates in SRC2v4-treated rhizospheres. Circles represent individual samples and bars reflect the mean of 4 biological replicates. Transcripts Per Million (TPM) for all genes were quantified by *Salmon*. **b)** Scatter plot of mean TPM/gene plotted in relation to the mean relative abundance of each SRC2v4 from a separate experiment (Fig. 2). Mean transcriptional activity significantly correlated with abundance (Spearman’s ρ = 0.70, p = 0.005).

**Figure S7.**
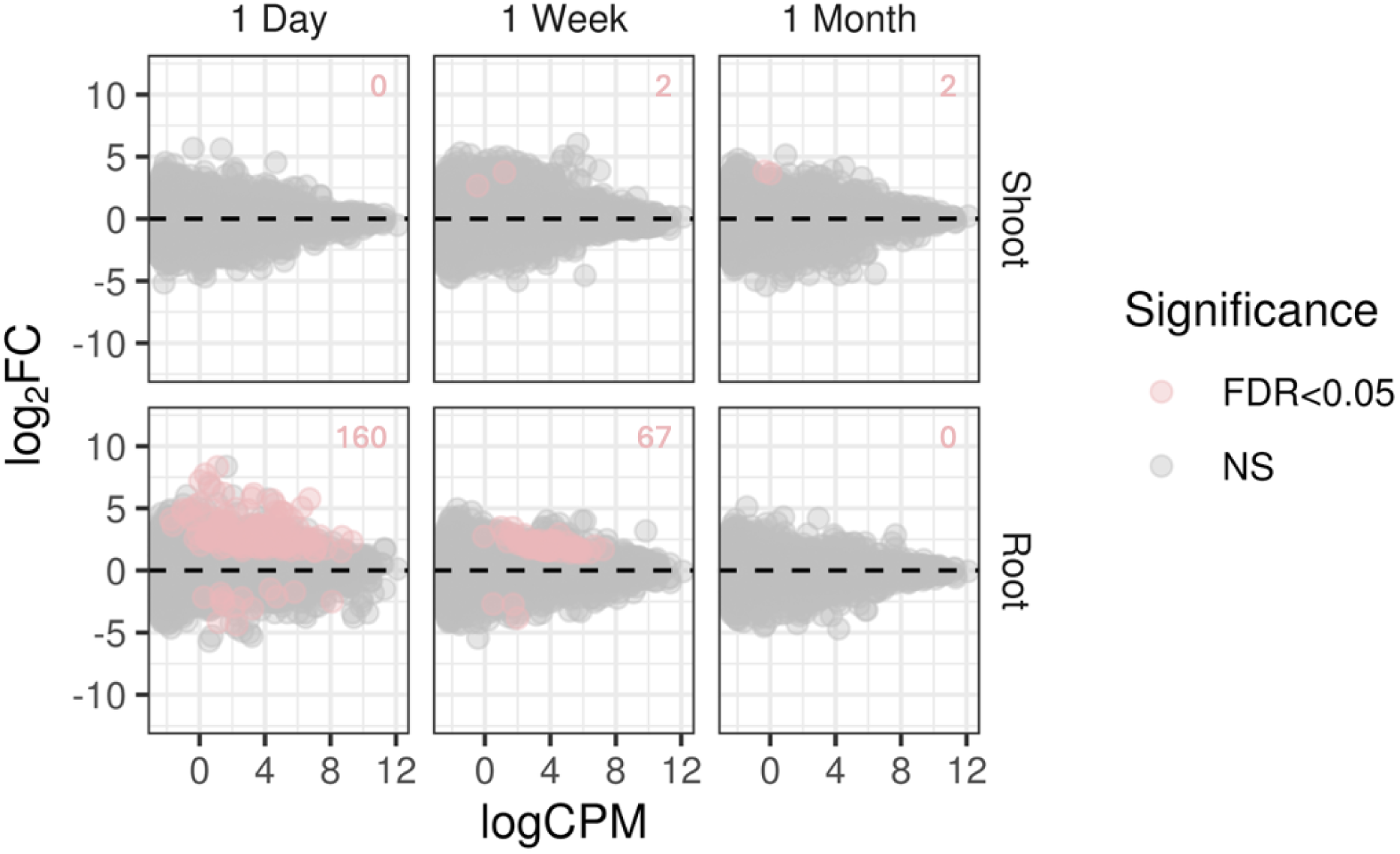
Sorghum host plants respond to HK treatment with an initial burst of regulation 1 day after treatment which tapers off in the following weeks and does not regulate genes in shoot tissue. Differentially expressed (DE) genes respond to HK treatment relative to Mock across three time points after treatment: 1 day, 1 week, and 1 month in both root (lower) and shoot (upper) tissue. Numbers in the upper right of each panel represent DE gene counts. Log_2_FC: log_2_-fold change (HK/Mock), logCPM: average log_2_counts per million across HK & Mock treatments, FDR<0.05: significantly differentially expressed genes after multiple testing correction, NS: not significant.

**Figure S8.**
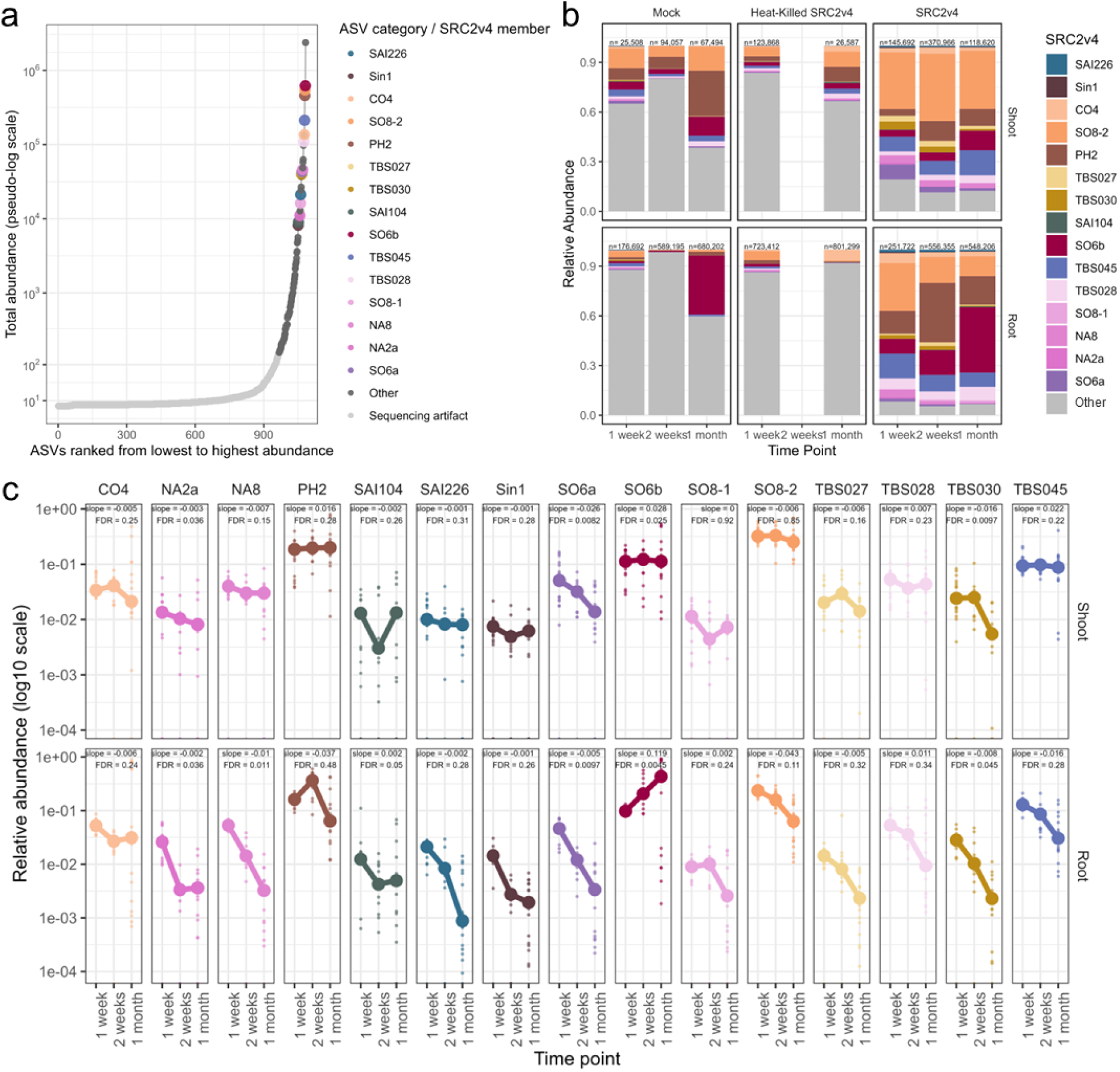
Temporal dynamics and relative abundance patterns of SRC2v4-associated ASVs in time-course experiment. **a)** Distribution of ASV abundances across the full dataset. ASVs are ranked from lowest to highest total abundance summed across all samples. Each point represents an individual ASV, colored by category: SRC2v4 members (colored individually by SRC2 member identity), environmental (non-SRC2) ASVs, or low-abundance ASVs classified as sequencing issues (total abundance <150). Abundances are displayed on a pseudo-logarithmic scale to accommodate the wide dynamic range of ASV abundances. **b)** Relative abundance of SRC2v4 members and environmental ASVs across time points and plant compartments. Relative abundances were calculated within samples and visualized to compare the contribution of SRC2v4-associated ASVs to the broader microbial community (total read count is shown above each bar). Environmental ASVs represent non-SRC2 taxa retained after filtering low-abundance without chloroplast and mitochondrial ASVs. (c) Temporal trends in relative abundance of individual SRC2v4 members across plant compartments. Points represent individual biological replicates, while lines and points indicate mean relative abundance at each time point. Linear models were fitted independently for each SRC2v4 member within each compartment to estimate abundance trends over time. Slopes and false discovery rate (FDR)-adjusted P values are shown for each SRC2v4 member. Relative abundances are plotted on a log10 scale with facet-specific y-axis scaling to facilitate comparison of temporal trends among SRC2v4 members spanning different abundance ranges.

**Figure S9.**
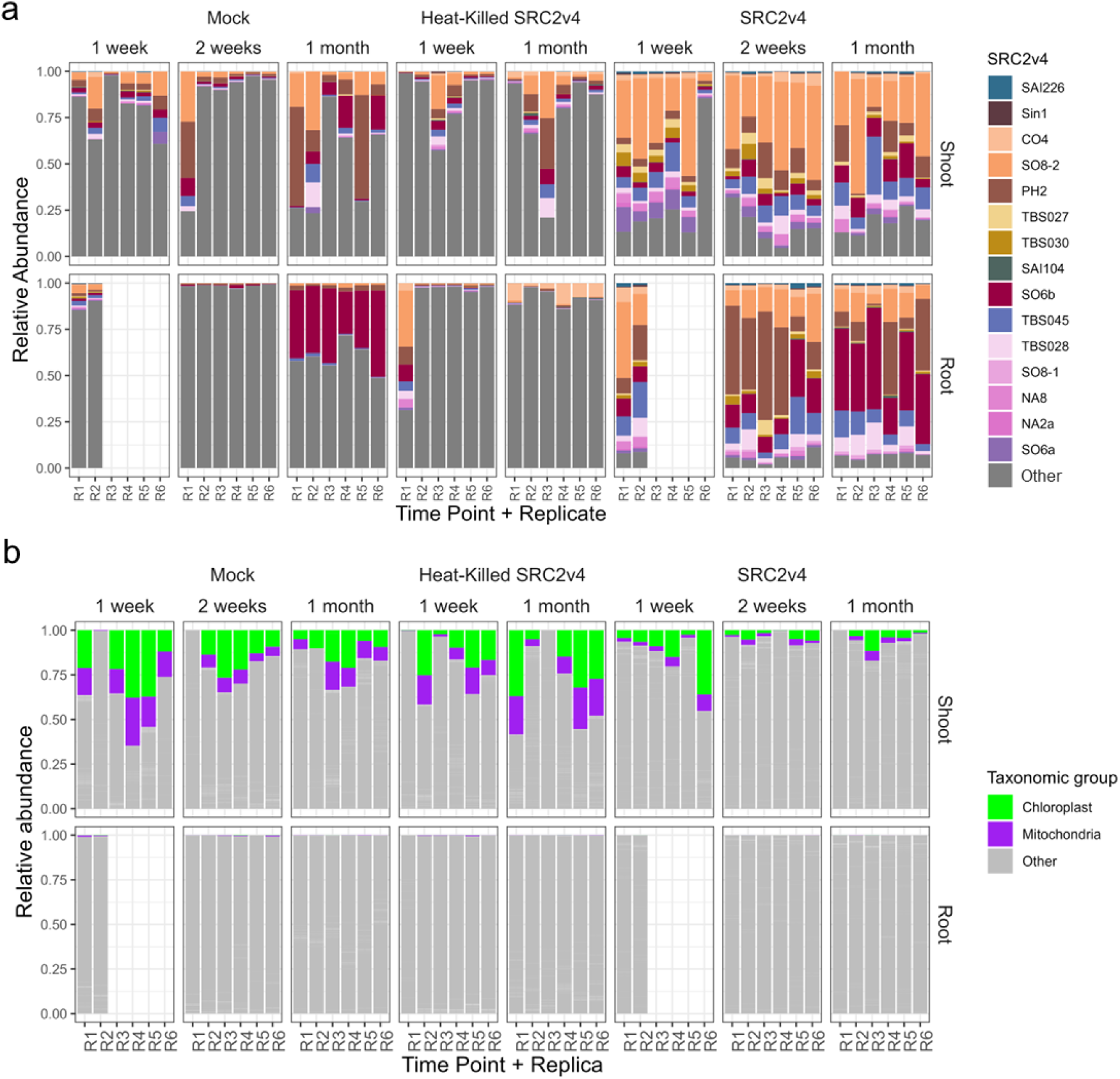
Relative abundance of SRC2v4 members and background ASVs across time course experiment. **a)** Relative abundance of SRC2v4 members across individual samples calculated from the full ASV table. Each bar represents one sample, and SRC2v4 members are colored individually. Low-abundance ASVs classified as sequencing issues and environmental (non-SRC2) ASVs are included in the total community in gray as Other. Relative abundances were calculated within each sample using unfiltered read counts. **b)** Relative abundance of chloroplast and mitochondria-associated ASVs across samples. Organelle-derived reads are shown explicitly to visualize their contribution to total sequencing output. All remaining non-SRC2, non-organelle ASVs are grouped and displayed in grey, representing the background microbial community.

**Figure S10.**
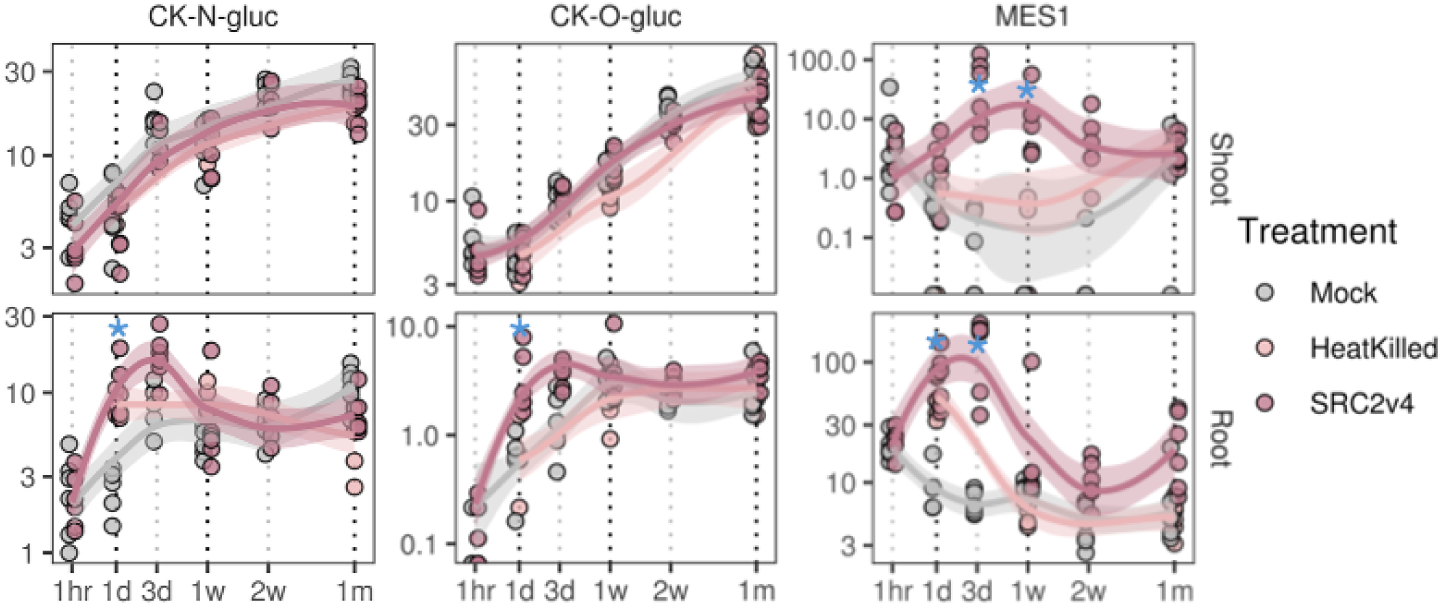
Spatiotemporal transcriptional regulation of key genes in SRC2v4 perception and plant response. Each gene is presented in two panels with treatment comparisons within each tissue type shoot (upper) and root (lower) across all time points with Loess fit trendlines by treatment. Hormone modulators include *Cytokinin-N-glucosyltransferase (*CK-N-Gluc), *Cytokinin-O-glucosyltransferase* (CK-O-Gluc), and *Methyl Esterase 1* (MES1). Blue stars indicate significant upregulation in SRC2v4 relative to Mock (Log_2_FoldChange > 1, FDR < 0.05), CPM = counts per million reads.

**Figure S11.**
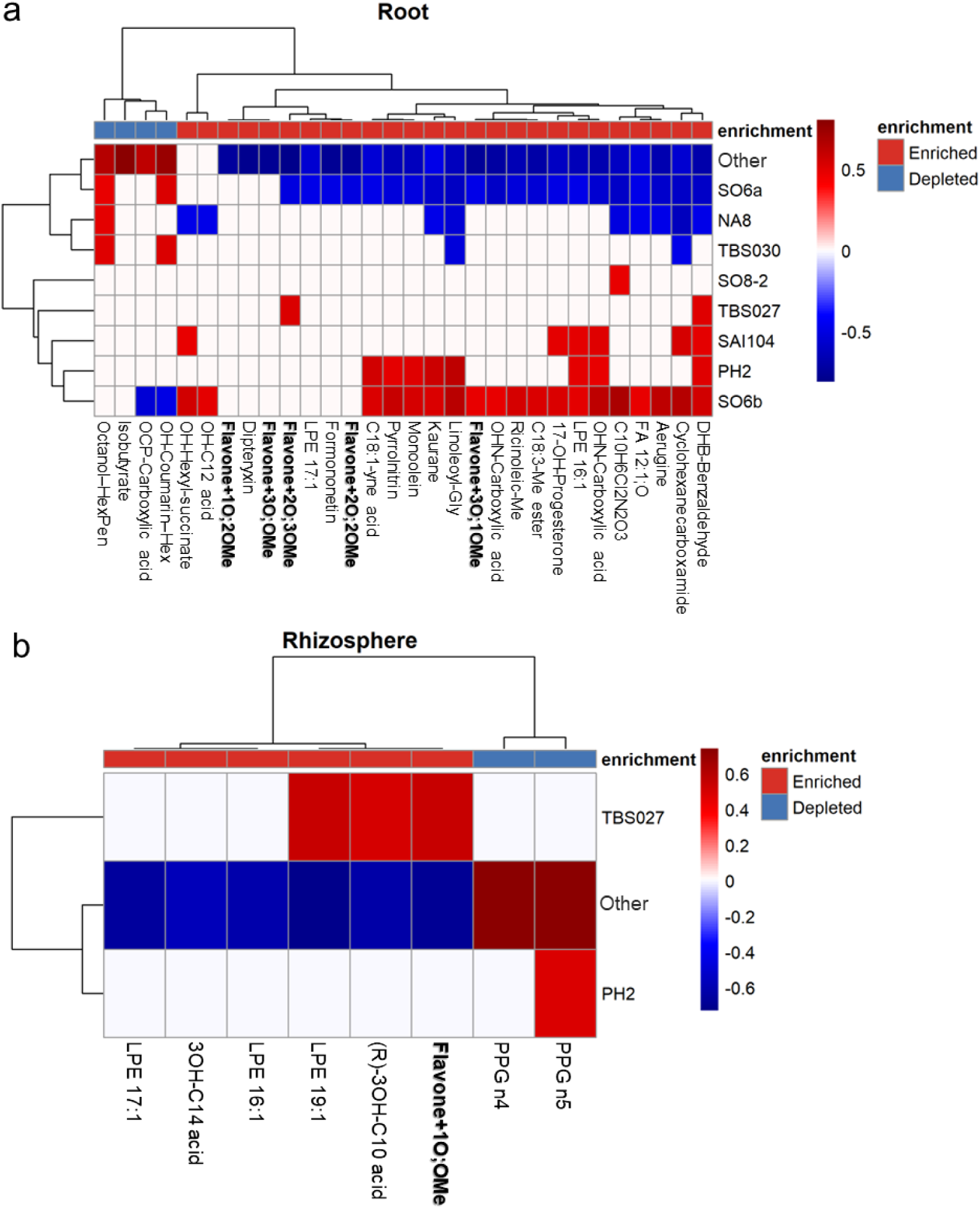
Associations between SRC2v4 members and root and rhizosphere metabolite profiles. Heatmaps show correlations between the relative abundances of individual SRC2v4 bacterial members derived from 16S rRNA amplicon sequencing and untargeted metabolite profiles in **a)** roots and **b)** rhizosphere samples. For the microbial data, SRC2v4 member abundances were calculated per sample, normalized to relative abundance, and transformed using a centered log-ratio (CLR) approach to account for compositionality. Metabolite intensities were log-transformed, z-score normalized across samples, and correlated with SRC2v4 CLR abundances using Spearman’s rank correlation. Only significant associations (FDR-adjusted P < 0.05) are shown and colors indicate the direction and strength of correlations (positive to negative). Rows correspond to individual SRC2v4 members, and columns correspond to annotated metabolites, clustered by correlation similarity. Metabolites are ordered according to enrichment or depletion in SRC2v4-inoculated samples relative to Mock controls. Correlations were calculated across Mock and SRC2v4 samples to capture treatment-associated variation. Metabolite abbreviations used in panel **a)**: 17-OH-Progesterone (17-hydroxyprogesterone); C18:3-Me (9(Z),11(E),13(E)-octadecatrienoic acid methyl ester); Ricinoleic-Me (ricinoleic acid methyl ester); OHN-carboxylic acid (2-(2-hydroxybutyl)-1,3-dimethyl-1,2,4a,5,6,7,8,8a-octahydronaphthalene-1-carboxylic acid); Flavone+3O;1OMe (flavone + 3O; 1OMe); Linoleoyl-Gly (linoleoyl glycerol); Kaurane (kaurane); Monoolein (monoolein); Pyrrolnitrin (pyrrolnitrin); C18:1-yne acid (octadecynoic acid); Flavone+2O;2OMe (flavone + 2O; 2OMe); Formononetin (formononetin); LPE17:1 (lysophosphatidylethanolamine 17:1); Flavone+2O;3OMe (flavone + 2O; 3OMe); Flavone+3O;OMe (flavone + 3O; OMe); Dipteryxin (dipteryxin); Flavone+1O;2OMe (flavone + 1O; 2OMe); OH-C12 acid (hydroxydodecanoate); OH-hexyl-succinate (2-(6-hydroxyhexyl)-3-methylenesuccinic acid); OH-coumarin-Hex (hydroxycoumarin + O-Hex); OCP-carboxylic acid (5a-hydroxy-1,7,7-trimethyl-1,2,3,3a,5a,6,7,8-octahydrocyclopenta[c]pentalene-4-carboxylic acid); Isobutyrate (isobutyric acid). Metabolite abbreviations used in panel **b)**: (R)-3OH-C10 acid ((R)-3-hydroxydecanoic acid); 3OH-C14 acid (3-hydroxytetradecanoic acid); LPE16:1 (lysophosphatidylethanolamine 16:1); LPE17:1 (lysophosphatidylethanolamine 17:1); LPE19:1 (lysophosphatidylethanolamine 19:1); PPG n4 (polypropylene glycol n4); PPG n5 (polypropylene glycol n5); Flavone+1O;OMe (flavone + 1O; OMe).

## References

Afrizal A, Jennings SAV, Hitch TCA, Riedel T, Basic M, Panyot A, Treichel N, Hager FT, Wong EO-Y, Wolter B, et al. 2022. Enhanced cultured diversity of the mouse gut microbiota enables custom-made synthetic communities. Cell Host & Microbe 30: 1630–1645.e25.

Appiah-Nkansah NB, Li J, Rooney W, Wang D. 2019. A review of sweet sorghum as a viable renewable bioenergy crop and its techno-economic analysis. Renewable energy 143: 1121–1132.

Banerjee S, Schlaeppi K, van der Heijden MGA. 2018. Keystone taxa as drivers of microbiome structure and functioning. Nature Reviews. Microbiology 16: 567–576.

Berihu M, Somera TS, Malik A, Medina S, Piombo E, Tal O, Cohen M, Ginatt A, Ofek-Lalzar M, Doron-Faigenboim A, et al. 2023. A framework for the targeted recruitment of crop-beneficial soil taxa based on network analysis of metagenomics data. Microbiome 11: 8.

Boller T, Felix G. 2009. A renaissance of elicitors: perception of microbe-associated molecular patterns and danger signals by pattern-recognition receptors. Annual review of plant biology 60: 379–406.

Bolyen E, Rideout JR, Dillon MR, Bokulich NA, Abnet CC, Al-Ghalith GA, Alexander H, Alm EJ, Arumugam M, Asnicar F, et al. 2019. Reproducible, interactive, scalable and extensible microbiome data science using QIIME 2. Nature biotechnology 37: 852–857.

Brenton ZW, Cooper EA, Myers MT, Boyles RE, Shakoor N, Zielinski KJ, Rauh BL, Bridges WC, Morris GP, Kresovich S. 2016. A genomic resource for the development, improvement, and exploitation of sorghum for bioenergy. Genetics 204: 21–33.

Bulgarelli D, Rott M, Schlaeppi K, Ver Loren van Themaat E, Ahmadinejad N, Assenza F, Rauf P, Huettel B, Reinhardt R, Schmelzer E, et al. 2012. Revealing structure and assembly cues for Arabidopsis root-inhabiting bacterial microbiota. Nature 488: 91–95.

Callahan BJ, McMurdie PJ, Rosen MJ, Han AW, Johnson AJA, Holmes SP. 2016. DADA2: High-resolution sample inference from Illumina amplicon data. Nature methods 13: 581–583.

Calviño M, Messing J. 2012. Sweet sorghum as a model system for bioenergy crops. Current opinion in biotechnology 23: 323–329.

Chen Y, Lun ATL, Smyth GK. 2016. From reads to genes to pathways: differential expression analysis of RNA-Seq experiments using Rsubread and the edgeR quasi-likelihood pipeline. F1000Research 5: 1438.

Chinchilla D, Zipfel C, Robatzek S, Kemmerling B, Nürnberger T, Jones JDG, Felix G, Boller T. 2007. A flagellin-induced complex of the receptor FLS2 and BAK1 initiates plant defence. Nature 448: 497–500.

Denoux C, Galletti R, Mammarella N, Gopalan S, Werck D, De Lorenzo G, Ferrari S, Ausubel FM, Dewdney J. 2008. Activation of defense response pathways by OGs and Flg22 elicitors in Arabidopsis seedlings. Molecular Plant 1: 423–445.

Díaz-Rodríguez AM, Parra Cota FI, Cira Chávez LA, García Ortega LF, Estrada Alvarado MI, Santoyo G, de Los Santos-Villalobos S. 2025. Microbial inoculants in sustainable agriculture: Advancements, challenges, and future directions. Plants 14: 191.

Dixon P. 2003. VEGAN, a package of R functions for community ecology. Journal of vegetation science: official organ of the International Association for Vegetation Science 14: 927–930.

Dobin A, Davis CA, Schlesinger F, Drenkow J, Zaleski C, Jha S, Batut P, Chaisson M, Gingeras TR. 2013. STAR: ultrafast universal RNA-seq aligner. Bioinformatics (Oxford, England) 29: 15–21.

Edwards J, Johnson C, Santos-Medellín C, Lurie E, Podishetty NK, Bhatnagar S, Eisen JA, Sundaresan V. 2015. Structure, variation, and assembly of the root-associated microbiomes of rice. Proceedings of the National Academy of Sciences of the United States of America 112: E911–20.

Emms DM, Kelly S. 2015. OrthoFinder: solving fundamental biases in whole genome comparisons dramatically improves orthogroup inference accuracy. Genome biology 16: 157.

Emms DM, Kelly S. 2019. OrthoFinder: phylogenetic orthology inference for comparative genomics. Genome biology 20: 238.

Finkel OM, Castrillo G, Herrera Paredes S, Salas González I, Dangl JL. 2017. Understanding and exploiting plant beneficial microbes. Current opinion in plant biology 38: 155–163.

Finkel OM, Salas-González I, Castrillo G, Conway JM, Law TF, Teixeira PJPL, Wilson ED, Fitzpatrick CR, Jones CD, Dangl JL. 2020. A single bacterial genus maintains root growth in a complex microbiome. Nature 587: 103–108.

Fonseca-García C, Pettinga D, Wilson A, Elmore JR, McClure R, Atim J, Pedraza J, Hutmacher R, Turumtay H, Tian Y, et al. 2024. Defined synthetic microbial communities colonize and benefit field-grown sorghum. The ISME journal 18.

Heese A, Hann DR, Gimenez-Ibanez S, Jones AME, He K, Li J, Schroeder JI, Peck SC, Rathjen JP. 2007. The receptor-like kinase SERK3/BAK1 is a central regulator of innate immunity in plants. Proceedings of the National Academy of Sciences of the United States of America 104: 12217–12222.

van der Heijden MGA, Bakker R, Verwaal J, Scheublin TR, Rutten M, van Logtestijn R, Staehelin C. 2006. Symbiotic bacteria as a determinant of plant community structure and plant productivity in dune grassland. FEMS Microbiology Ecology 56: 178–187.

He D, Singh SK, Peng L, Kaushal R, Vílchez JI, Shao C, Wu X, Zheng S, Morcillo RJL, Paré PW, et al. 2022. Flavonoid-attracted Aeromonas sp. from the Arabidopsis root microbiome enhances plant dehydration resistance. The ISME journal 16: 2622–2632.

He Z, Wang ZY, Li J, Zhu Q, Lamb C, Ronald P, Chory J. 2000. Perception of brassinosteroids by the extracellular domain of the receptor kinase BRI1. Science (New York, N.Y.) 288: 2360–2363.

Jeong SY, Park CH, Kim M-K, Nam SJ, Hong J, Kim S-K. 2012. Effect of lysophosphatidylethanolamine and brassinosteroids on development of Arabidopsis roots. Journal of Plant Biology 55: 178–184.

Karp RM. 1972. Reducibility among Combinatorial Problems. In: Complexity of Computer Computations. Boston, MA: Springer US, 85–103.

Kumar GA, Kumar S, Bhardwaj R, Swapnil P, Meena M, Seth CS, Yadav A. 2023. Recent advancements in multifaceted roles of flavonoids in plant-rhizomicrobiome interactions. Frontiers in plant science 14: 1297706.

Kwak M-J, Kong HG, Choi K, Kwon S-K, Song JY, Lee J, Lee PA, Choi SY, Seo M, Lee HJ, et al. 2018. Rhizosphere microbiome structure alters to enable wilt resistance in tomato. Nature Biotechnology 36: 1100–1109.

Lee MD. 2019. GToTree: a user-friendly workflow for phylogenomics. Bioinformatics 35: 4162–4164.

Levy A, Salas Gonzalez I, Mittelviefhaus M, Clingenpeel S, Herrera Paredes S, Miao J, Wang K, Devescovi G, Stillman K, Monteiro F, et al. 2017. Genomic features of bacterial adaptation to plants. Nature genetics 50: 138–150.

Li J, Chory J. 1997. A putative leucine-rich repeat receptor kinase involved in brassinosteroid signal transduction. Cell 90: 929–938.

Li J, Wen J, Lease KA, Doke JT, Tax FE, Walker JC. 2002. BAK1, an Arabidopsis LRR receptor-like protein kinase, interacts with BRI1 and modulates brassinosteroid signaling. Cell 110: 213–222.

Lundberg DS, Lebeis SL, Paredes SH, Yourstone S, Gehring J, Malfatti S, Tremblay J, Engelbrektson A, Kunin V, Rio TG del, et al. 2012. Defining the core Arabidopsis thaliana root microbiome. Nature 488: 86–90.

Macho AP, Zipfel C. 2014. Plant PRRs and the activation of innate immune signaling. Molecular cell 54: 263–272.

Mahon EL, de Vries L, Jang S-K, Middar S, Kim H, Unda F, Ralph J, Mansfield SD. 2022. Exogenous chalcone synthase expression in developing poplar xylem incorporates naringenin into lignins. Plant physiology 188: 984–996.

Martins SJ, Pasche J, Silva HAO, Selten G, Savastano N, Abreu LM, Bais HP, Garrett KA, Kraisitudomsook N, Pieterse CMJ, et al. 2023. The use of synthetic microbial communities to improve plant health. Phytopathology 113: 1369–1379.

Montoya M, Durán-Wendt D, Garrido-Sanz D, Carrera-Ruiz L, Vázquez-Arias D, Redondo-Nieto M, Martín M, Rivilla R. 2025. Functional characterization of a synthetic bacterial community (SynCom) and its impact on gene expression and growth promotion in tomato. Agronomy (Basel, Switzerland) 15: 1794.

Mueller UG, Sachs JL. 2015. Engineering microbiomes to improve plant and animal health. Trends in microbiology 23: 606–617.

Nam KH, Li J. 2002. BRI1/BAK1, a receptor kinase pair mediating brassinosteroid signaling. Cell 110: 203–212.

Niu B, Paulson JN, Zheng X, Kolter R. 2017. Simplified and representative bacterial community of maize roots. Proceedings of the National Academy of Sciences of the United States of America 114: E2450–E2459.

Paasch BC, Sohrabi R, Kremer JM, Nomura K, Cheng YT, Martz J, Kvitko B, Tiedje JM, He SY. 2023. A critical role of a eubiotic microbiota in gating proper immunocompetence in Arabidopsis. Nature Plants 9: 1468–1480.

Pang Z, Lu Y, Zhou G, Hui F, Xu L, Viau C, Spigelman AF, MacDonald PE, Wishart DS, Li S, et al. 2024. MetaboAnalyst 6.0: towards a unified platform for metabolomics data processing, analysis and interpretation. Nucleic acids research 52: W398–W406.

Patro R, Duggal G, Love MI, Irizarry RA, Kingsford C. 2017. Salmon provides fast and bias-aware quantification of transcript expression. Nature Methods 14: 417–419.

Peters NK, Frost JW, Long SR. 1986. A plant flavone, luteolin, induces expression of Rhizobium meliloti nodulation genes. Science (New York, N.Y.) 233: 977–980.

Philippot L, Raaijmakers JM, Lemanceau P, van der Putten WH. 2013. Going back to the roots: the microbial ecology of the rhizosphere. Nature reviews. Microbiology 11: 789–799.

Pineda Rodo A, Brugière N, Vankova R, Malbeck J, Olson JM, Haines SC, Martin RC, Habben JE, Mok DWS, Mok MC. 2008. Over-expression of a zeatin O-glucosylation gene in maize leads to growth retardation and tasselseed formation. Journal of Experimental Botany 59: 2673–2686.

R Core Team. 2023. R: A Language and Environment for Statistical Computing. Vienna, Austria: R Foundation for Statistical Computing.

Redmond JW, Batley M, Djordjevic MA, Innes RW, Kuempel PL, Rolfe BG. 1986. Flavones induce expression of nodulation genes in Rhizobium. Nature 323: 632–635.

Robinson MD, McCarthy DJ, Smyth GK. 2010. edgeR: a Bioconductor package for differential expression analysis of digital gene expression data. Bioinformatics 26: 139–140.

Shi J, Yan X, Sun T, Shen Y, Shi Q, Wang W, Bao M, Luo H, Nian F, Ning G. 2022. Homeostatic regulation of flavonoid and lignin biosynthesis in phenylpropanoid pathway of transgenic tobacco. Gene 809: 146017.

Shumilina J, Soboleva A, Abakumov E, Shtark OY, Zhukov VA, Frolov A. 2023. Signaling in legume-rhizobia symbiosis. International journal of molecular sciences 24: 17397.

Šmehilová M, Dobrůšková J, Novák O, Takáč T, Galuszka P. 2016. Cytokinin-specific glycosyltransferases possess different roles in cytokinin homeostasis maintenance. Frontiers in Plant Science 7: 1264.

de Souza RSC, Armanhi JSL, Arruda P. 2020. From microbiome to traits: Designing synthetic microbial communities for improved crop resiliency. Frontiers in plant science 11: 1179.

de Souza RSC, Armanhi JSL, Damasceno N de B, Imperial J, Arruda P. 2019. Genome sequences of a plant beneficial synthetic bacterial community reveal genetic features for successful plant colonization. Frontiers in microbiology 10: 1779.

Stringlis IA, Proietti S, Hickman R, Van Verk MC, Zamioudis C, Pieterse CMJ. 2018. Root transcriptional dynamics induced by beneficial rhizobacteria and microbial immune elicitors reveal signatures of adaptation to mutualists. The Plant Journal: For Cell and Molecular Biology 93: 166–180.

Teixeira PJPL, Colaianni NR, Law TF, Conway JM, Gilbert S, Li H, Salas-González I, Panda D, Del Risco NM, Finkel OM, et al. 2021. Specific modulation of the root immune system by a community of commensal bacteria. Proceedings of the National Academy of Sciences of the United States of America 118: e2100678118.

Trivedi P, Leach JE, Tringe SG, Sa T, Singh BK. 2020. Plant-microbiome interactions: from community assembly to plant health. Nature reviews. Microbiology 18: 607–621.

Tzipilevich E, Russ D, Dangl JL, Benfey PN. 2021. Plant immune system activation is necessary for efficient root colonization by auxin-secreting beneficial bacteria. Cell Host & Microbe 29: 1507–1520.e4.

Varoquaux N, Cole B, Gao C, Pierroz G, Baker CR, Patel D, Madera M, Jeffers T, Hollingsworth J, Sievert J, et al. 2019. Transcriptomic analysis of field-droughted sorghum from seedling to maturity reveals biotic and metabolic responses. Proceedings of the National Academy of Sciences of the United States of America 116: 27124–27132.

Weston LA, Alsaadawi IS, Baerson SR. 2013. Sorghum allelopathy--from ecosystem to molecule. Journal of Chemical Ecology 39: 142–153.

Wu Y, Ma Y, Wang W, Zhang S, Wu W. 2025. Mastering the balance: BAK1’s dual roles in steering plant growth and immunity. Horticulture Research 12: uhaf206.

Xu X, Dinesen C, Pioppi A, Kovács ÁT, Lozano-Andrade CN. 2025. Composing a microbial symphony: synthetic communities for promoting plant growth. Trends in microbiology 33: 738–751.

Xu L, Naylor D, Dong Z, Simmons T, Pierroz G, Hixson KK, Kim Y-M, Zink EM, Engbrecht KM, Wang Y, et al. 2018. Drought delays development of the sorghum root microbiome and enriches for monoderm bacteria. Proceedings of the National Academy of Sciences of the United States of America 115: E4284–E4293.

Yang X, Owens TG, Scheffler BE, Weston LA. 2004. Manipulation of root hair development and sorgoleone production in sorghum seedlings. Journal of Chemical Ecology 30: 199–213.

Yang Y, Xu R, Ma C-J, Vlot AC, Klessig DF, Pichersky E. 2008. Inactive methyl indole-3-acetic acid ester can be hydrolyzed and activated by several esterases belonging to the AtMES esterase family of Arabidopsis. Plant physiology 147: 1034–1045.

Yan D, Tajima H, Cline LC, Fong RY, Ottaviani JI, Shapiro H-Y, Blumwald E. 2022. Genetic modification of flavone biosynthesis in rice enhances biofilm formation of soil diazotrophic bacteria and biological nitrogen fixation. Plant Biotechnology Journal 20: 2135–2148.

Yu P, He X, Baer M, Beirinckx S, Tian T, Moya YAT, Zhang X, Deichmann M, Frey FP, Bresgen V, et al. 2021. Plant flavones enrich rhizosphere Oxalobacteraceae to improve maize performance under nitrogen deprivation. Nature plants 7: 481–499.

Zhuang L, Li Y, Wang Z, Yu Y, Zhang N, Yang C, Zeng Q, Wang Q. 2021. Synthetic community with six Pseudomonas strains screened from garlic rhizosphere microbiome promotes plant growth. Microbial Biotechnology 14: 488–502.

